# Pervasive and programmed nucleosome distortion patterns on single mammalian chromatin fibers

**DOI:** 10.1101/2025.01.17.633622

**Authors:** Marty G Yang, Hannah J Richter, Simai Wang, Colin P McNally, Nicole Harris, Simaron Dhillon, Michela Maresca, Elzo de Wit, Holger Willenbring, Jacquelyn Maher, Hani Goodarzi, Vijay Ramani

## Abstract

We present a genome-scale method to map the single-molecule co-occupancy of structurally distinct nucleosomes, subnucleosomes, and other protein-DNA interactions via long-read high-resolution adenine methyltransferase footprinting. <u>I</u>teratively <u>D</u>efined <u>L</u>engths of Inaccessibility (IDLI) classifies nucleosomes on the basis of shared patterns of intranucleosomal accessibility, into: i.) minimally-accessible chromatosomes; ii.) octasomes with stereotyped DNA accessibility from superhelical locations (SHLs) ±1 through ±7; iii.) highly-accessible unwrapped nucleosomes; and iv.) subnucleosomal species, such as hexasomes, tetrasomes, and other short DNA protections. Applying IDLI to mouse embryonic stem cell (mESC) chromatin, we discover widespread nucleosomal distortion on individual mammalian chromatin fibers, with >85% of nucleosomes surveyed displaying degrees of intranucleosomally accessible DNA. We observe epigenomic-domain-specific patterns of distorted nucleosome co-occupancy and positioning, including at enhancers, promoters, and mouse satellite repeat sequences. Nucleosome distortion is programmed by the presence of bound transcription factors (TFs) at cognate motifs; occupied TF binding sites are differentially decorated by distorted nucleosomes compared to unbound sites, and degradation experiments establish direct roles for TFs in structuring binding-site proximal nucleosomes. Finally, we apply IDLI in the context of primary mouse hepatocytes, observing evidence for pervasive nucleosomal distortion *in vivo.* Further genetic experiments reveal a role for the hepatocyte master regulator FOXA2 in directly impacting nucleosome distortion at hepatocyte-specific regulatory elements *in vivo*. Our work suggests extreme—but regulated—plasticity in nucleosomal DNA accessibility at the single-molecule level. Further, our study offers an essential new framework to model transcription factor binding, nucleosome remodeling, and cell-type specific gene regulation across biological contexts.

## INTRODUCTION

Regulated DNA accessibility governs cell-type specific transcription. DNA accessibility is dictated in part by the octameric nucleosome, which compacts ∼147 basepairs (bp) of DNA around two dimers of histone proteins H2A and H2B flanking a tetramer of histone proteins H3 and H4. Our canonical understanding of how chromatin accessibility regulates specificity of protein-DNA interactions is simple: nucleosomes and positive transcriptional regulatory factors compete with one another for DNA binding, particularly at active *cis*-regulatory elements^1^. Thus, nucleosome occupancy negatively regulates transcription, and multiple essential coactivators act by modulating nucleosome occupancy directly via chromatin remodeling.

Work over the last three decades has challenged the simplicity of this model. The synthesis of genome-scale mapping techniques and high-resolution structural biology has revealed ‘altered’ nucleosomal structures across eukaryotic epigenomes, including hexasomes (*i.e.* nucleosomes missing a single H2A/H2B dimer)^2–4^, overlapping dinucleosomes (*i.e.* hexasome-octasome complexes)^5–7^, and other noncanonical histone-containing complexes^8^. Work from the groups of Widom and others has illustrated that primary DNA sequence can guide occupancy and modulate the stability of nucleosome-wrapped DNA^9–11^. ATP-dependent chromatin remodeling can rely on structural distortion of nucleosomes (for the imitation switch [ISWI] remodeler SNF2h^12^), or the presence of hexasomes (for INO80^13,14^). Thus, dynamic nucleosome composition and structure are physiologically-relevant: nuclear regulatory processes—including ATP-dependent chromatin remodeling, transcription, replication, and histone chaperone activity—generate these species and render them important marks of genomic activity. Perhaps most importantly, all of these species must ultimately compete with essential sequence-specific transcriptional factors (TFs) to determine cell-type-specific chromatin accessibility and gene transcription^15^. Holistically understanding how nucleosomes are structured and positioned with respect to TFs necessitates high-resolution *in vivo* measurement of nucleosomal structures at TF binding sites across diverse contexts.

Technical limitations have confounded accurate assessment of canonical and noncanonical nucleosome structures *in vivo*. Highly resolved *in situ* imaging techniques (*e.g.* [cryo]electron tomography^16,17^; chromatin expansion microscopy^18^) are blind to DNA sequence, while standard genomic mapping techniques rely on digestion of the chromatin fiber^3,4,19,20^. Digestion is especially problematic in the context of mapping subnucleosomal particles: a hexasome-sized nucleolytic fragment, for instance, could either derive from a true hexasome, from a remodeled nucleosome with increased DNA accessibility between a single H2A/H2B dimer and the H3/H4 tetramer, or from transient unwrapping of DNA at nucleosomal entry / exit sites (*i.e.* nucleosomal breathing)^21,22^. Short read single-molecule methyltransferase (MTase) footprinting methods (*e.g.* NoME-seq^23^; MAPit^24^; dSMF^25^) solve some of these issues but are themselves limited by i.) the maximum read-length (600 bp) possible on an Illumina sequencer, and ii.) use of MTases with specificity for GpC or CpG dinucleotides. Single-molecule long-read footprinting methods that use promiscuous adenine MTases (*e.g.* the <u>s</u>ingle-molecule <u>a</u>denine <u>m</u>ethylated <u>o</u>ligonucleosome <u>s</u>equencing <u>a</u>ssay [SAMOSA]^26,27^; Fiber-seq^28^) theoretically solve many of these limitations, but to date have not been applied towards systematically interrogating noncanonical nucleosome structure.

Here, we build on computational methods previously developed by our group^29^ to nondestructively resolve—for the first time—the subunit architecture of chromatin fibers down to individual H2A/H2B dimers and H3/H4 tetramers. Harnessing this ability to map structurally distinct classes of nucleosomes on chromatin fibers genome-wide *in vivo*, we make a startling discovery: pervasive, but regulated patterns of ‘distorted’ nucleosomes with varying degrees of accessible nucleosomal DNA, across both active and inactive mammalian epigenomic regions. Acute depletion of transcription factors (CTCF; SOX2) in mouse embryonic stem cells (mESCs) reveals distinct modes by which essential trans-acting factors distort and position nucleosomes on chromatin fibers. Extension of our approach to data generated in primary mouse hepatocytes reveals that widespread nucleosome distortion occurs *in vivo*. By genetically perturbing the master regulatory factor FOXA2 in hepatocytes *in vivo*, we demonstrate a direct role for this ‘pioneer’ factor in positioning and distorting nucleosomes at FOXA2-bound regulatory regions. Taken as a whole, our study provides a sequence-resolved foundation for quantifying and studying causes and consequences of nucleosome structural heterogeneity genome-wide.

## RESULTS

### Footprinting the subunit architecture of the nucleosome core particle on individual chromatin fibers

We previously introduced the <u>s</u>ingle-molecule <u>a</u>denine <u>m</u>ethylated <u>o</u>ligonucleosome <u>s</u>equencing <u>a</u>ssay (SAMOSA), which combines genome-wide chromatin footprinting by nonspecific adenine MTase EcoGII with native detection of methylated adenine via long-read PacBio sequencing^26^. Inferring protein-DNA contacts from EcoGII methylation is nontrivial, and we later introduced a neural network – hidden markov model (NN-HMM) that explicitly accounts for: i.) sequence biases of the EcoGII enzyme, and ii.) sequence biases influencing the kinetics of PacBio sequencing, to infer nucleosome ‘footprint’ positions along sequenced DNA molecules^29^. Our original approach – which has successfully mapped chromatin *in vivo*^27^, *in vitro*^29,30^, and in the context of DNA replication^31^ – was calibrated to detect mononucleosomes, the predominant feature of chromatin.

We hypothesized that our model – which uses an NN to define emission probabilities and requires only a single, user-defined parameter (the transition probability; hereafter, ‘*t*’) within a two-state HMM – might be able to detect the subunit structure of mononucleosomes (**Figure 1A**). To test this hypothesis, we collated a mixture of published and unpublished SAMOSA datasets for the E14 mESC line (44 sequencing libraries to yield 88.87 Gb of sequenced mESC DNA; library metadata summarized in **Supplementary Table 1**), and called footprints varying the *t* parameter (which represents the likelihood of switching between hidden ‘inaccessible’ and ‘accessible’ model states). Plotting the distributions of called footprint sizes across a wide range of *t* values confirmed that increasing *t* resulted in successively smaller footprints calls (hereafter referred to as ‘subfootprints’; **Figure 1B**). We reasoned that if subfootprints were biologically ‘patterned,’ then due to the two-fold symmetry of the nucleosome about the dyad, accessibility surrounding subfootprints should symmetrically shift when plotted about the center of the originating footprint (due to nucleosomal ‘breathing’ at the entry / exit site^32^); further, we reasoned that these footprints, when plotted as footprint size versus midpoint position ‘horizon plots’ (analogous to V-plots for micrococcal nuclease sequencing [MNase-seq] data^19^), would reflect the composition of the nucleosome, including H2A/H2B dimers, H3/H4 tetramer, hexasomes, and “breathing” nucleosomes (**Figure 1C-D**).

**Figure 1:**
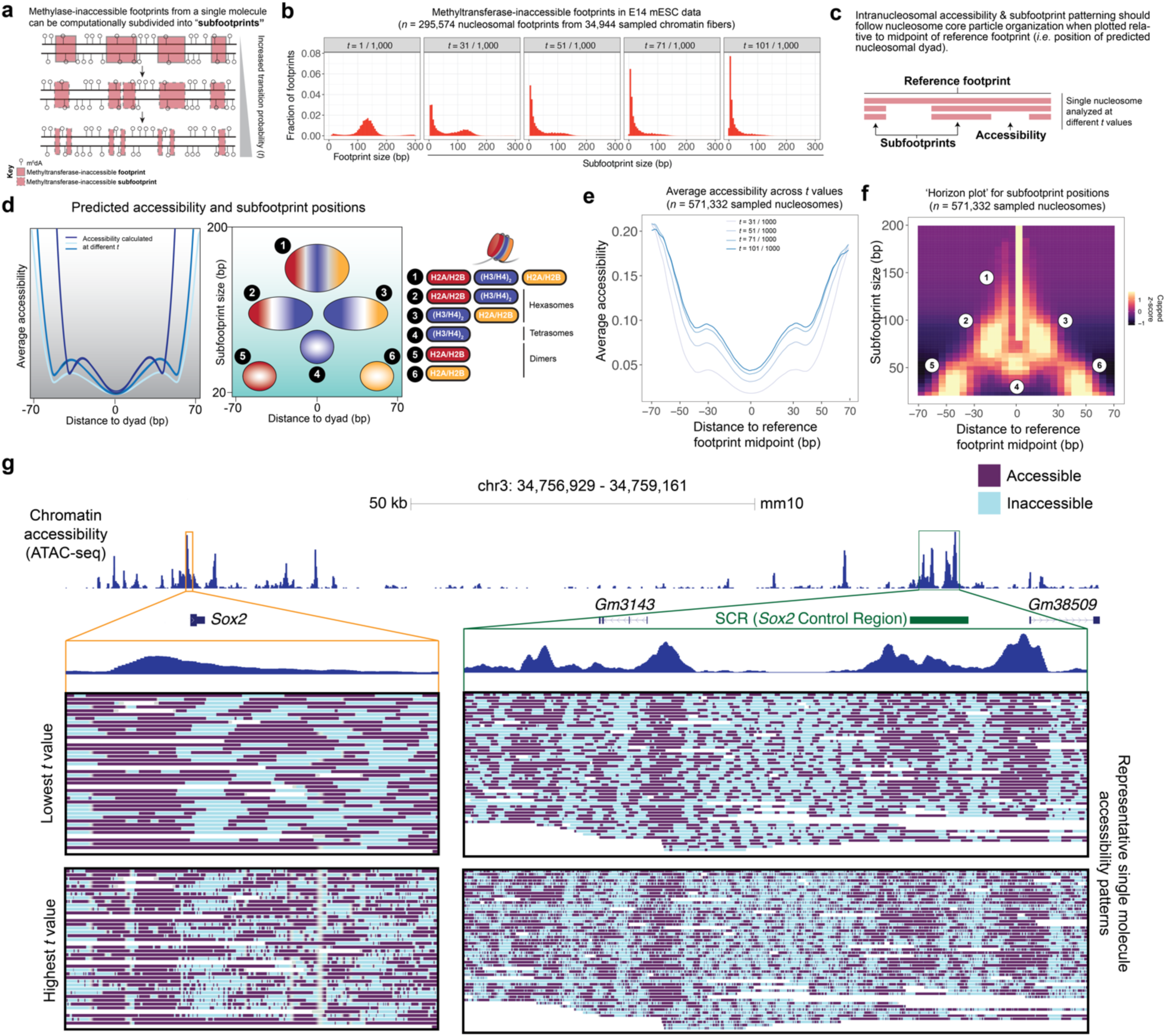
Computational framework for defining subunit architecture and organization of nucleosome core particles from *in vivo* long-read methyltransferase footprinting data. **A.)** Sequencing reads from <u>s</u>ingle-molecule <u>a</u>denine <u>m</u>ethylated <u>o</u>ligonucleosome <u>s</u>equencing <u>a</u>ssay [SAMOSA] data are analyzed with a two-state hidden Markov model (HMM) to define contiguous stretches of methylase-inaccessible nucleotides (‘footprints’) from EcoGII-treated chromatin. As shown in this schematic, individual footprints are subdivided into shorter ‘subfootprints’ at higher transition probabilities (‘*t*’; the likelihood of switching between the ‘inaccessible’ and ‘accessible’ model states). **B.)** Distributions of footprint and subfootprint sizes from randomly sampled footprints from E14 mESCs across a range of *t* values. Shown on the left is the footprint size distribution from the transition probability (*t* = 1 / 1,000) used in our prior work, calibrated to principally detect mononucleosome-sized footprints. Increasing *t* values results in a greater proportion of progressively shorter footprints from an identical set of chromatin fibers. **C.)** We named this computational framework wherein intra-nucleosomal accessibility within single footprints are systematically analyzed across a range of *t* values: <u>I</u>teratively <u>D</u>efined <u>L</u>engths of <u>I</u>naccessibility (or IDLI, for short). We hypothesized that by tuning the transition probability, we should be able to recapitulate intra-nucleosome accessibility and subfootprint patterns reflective of the organization of the nucleosome core particle. **D.)** Predicted outcomes for nucleosomal footprints in E14 mESC data analyzed with IDLI. **Left:** Mean accessibility should be patterned symmetrically with respect to the footprint midpoints (*i.e.* predicted nucleosomal dyad positions) at different *t* values. **Right:** Predicted sizes and positions of subfootprints (plotted relative to footprint midpoints) should reflect the subunit architecture of the octameric nucleosome, including: (1) histone octamers, (2-3) hexasomes (lacking an H2A-H2B dimer), (4) H3-H4 tetramers, and (5-6) H2A-H2B dimers. **E.)** Measured mean accessibility with respect to reference (*i.e.* from *t* = 1 / 1,000) footprint midpoints reveals expected symmetric pattern about predicted dyad positions. **F.)** ‘Horizon plot’ visualization (with respect to footprint midpoints) captures enrichment of previously-noted subfootprint sizes and positions, including octameric nucleosomes that exhibit spontaneous ‘unspooling’ of DNA from entry / exit sites (Feature 1; vertical line centered at footprint midpoint). **G.)** Single-molecule accessibility patterns (EcoGII-accessible and -inaccessible DNA in purple and teal, respectively) at *Sox2* promoter (left; depicted in orange) and *Sox2* Control Region [SCR] (right; depicted in green). Data from representative identical single EcoGII-footprinted fibers are shown for the lowest (*t* = 1 / 1,000; top) and highest (*t* = 101 / 1,000; bottom) transition probabilities used in IDLI. Bulk-average chromatin accessibility data (from ATAC-seq) in mESCs are also shown across the entire locus (top) and at specific regulatory loci (bottom).

Visualization of data in both ways confirmed that our analysis of subfootprints captured the expected intranucleosomal accessibility and patterns of protein-DNA contacts from nucleosomes *in vivo*. At four different *t* values, accessibility demonstrated a symmetric pattern about called footprint midpoints (**Figure 1E**). Most strikingly, horizon plot visualization of subfootprints revealed clear signal enrichment at precise positions (**Figure 1F**), indicative of subfootprinting of breathing octasomes (vertical line; feature 1), hexasomes (feature 2), tetramers (feature 3), and dimers (feature 4). Importantly, this result was quantitatively robust across multiple biological replicates (**Extended Data Figure 1A-B**; median correlation ± median absolute deviation [MAD] = 0.906 ± 0.010), and robust to per-sample methylation amount (measured through the mononucleosome-dinucleosome ratio [MDR]^33^; see **Methods**; **Extended Data Figure 1C**). Finally, accessibility and footprint / subfootprint positions on individual fibers were easily visualized along with bulk ATAC-seq data^34^ at the *Sox2* locus (**Figure 1G**; **Extended Data Figure 1D**). Taken together, these results demonstrate our ability to footprint the subunit architecture of the nucleosome on individual chromatin fibers in a mammalian epigenome. Hereafter, we refer to this computational strategy—dependent on iteratively scoring footprinted molecules—as Iteratively <u>D</u>efined <u>L</u>engths of <u>I</u>naccessibility, or IDLI.

### The mammalian epigenome is composed of a constellation of structurally distinct nucleosomes

We next sought to evaluate structural heterogeneity as defined by IDLI, at the resolution of individual nucleosome footprints. We devised a classification approach (**Extended Data Figure 2A**) that leverages the Leiden community detection^35^ and UMAP visualization^36^ algorithms to classify and visualize structurally distinct nucleosome footprints defined by IDLI. We applied this approach to 795,765 sampled nucleosomes from 145,065 sampled E14 mESC chromatin fibers falling within 8 distinct epigenomic regions, including both mappable domains of specific post-translationally modified histones and typically ‘unmappable’ domains comprised of low-complexity repeat sequence (mouse major satellite, minor satellite, and telomeric sequence). We further binned clusters into ‘groups’ through hierarchical clustering (**Extended Data Figure 2B**) and examined the underlying properties of each cluster of nucleosomes. Specifically, we examined three properties (**Extended Data Figure 2C**): i.) per-cluster footprint sizes; ii.) per-cluster horizon plots; and iii.) average accessibility of footprints at the most permissive calculated *t* value used for clustering (*i.e., t* = 0.071). Our joint analysis and visualization pipeline (UMAP colored by cluster shown in **Figure 2A**) resulted in 14 distinct structural classes (*i.e.* ‘nucleosome types;’ **Figure 2B**), which fell into seven larger groups.

**Figure 2:**
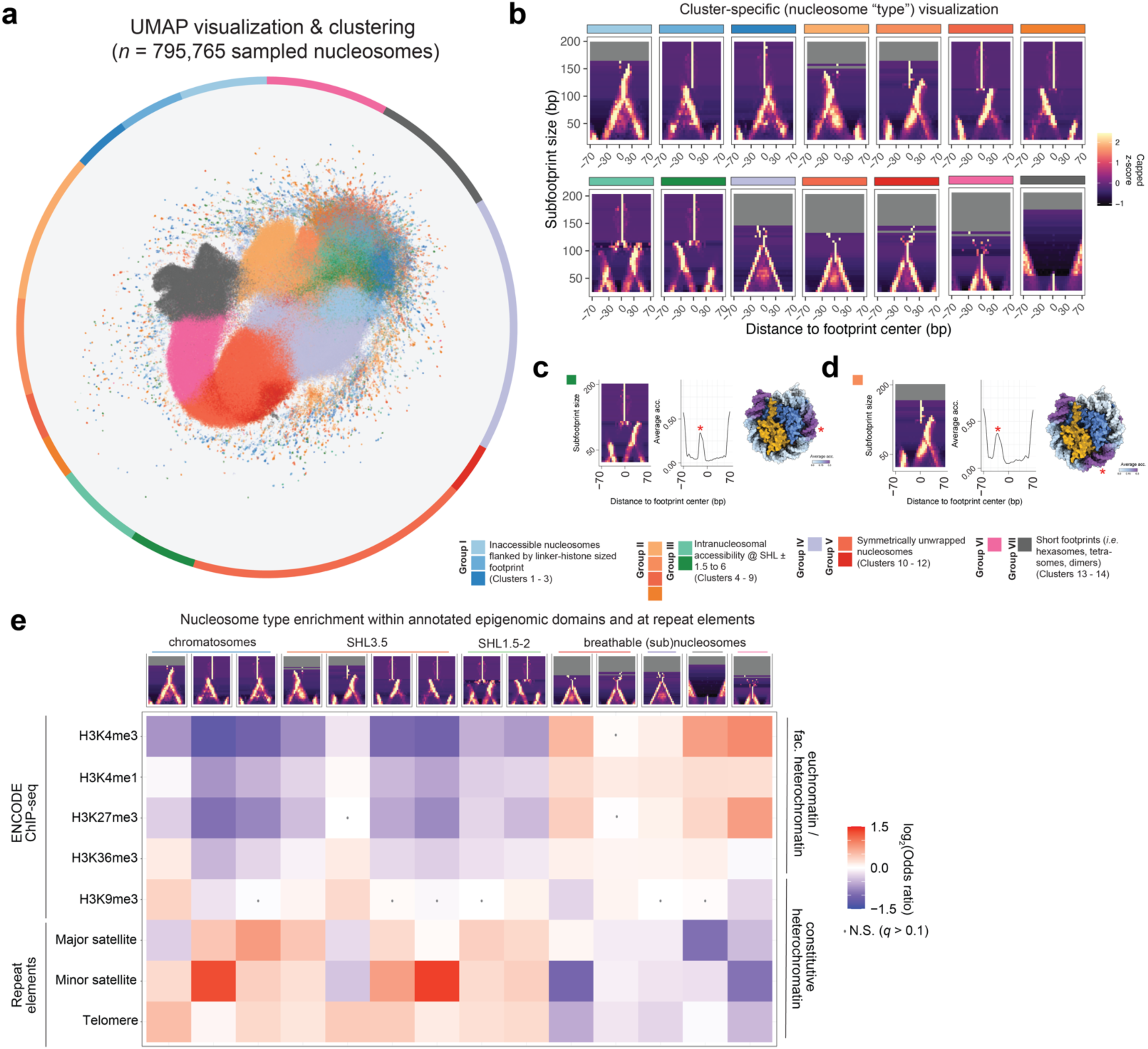
Structurally distinct nucleosomes are enriched in an epigenomic domain- and repeat-sequence-specific manner. **A.)** UMAP visualization of *n* = 795,765 footprints within ±500 bp of midpoints of histone post-translational modification-defined epigenomic domains (in mESCs) or ‘unmappable’ repeat elements. Individual footprints within UMAP are colored based on Leiden-clustering assignments of their intra-nucleosomal accessibility (*i.e.* distortion patterns). Proportions of each cluster are depicted as a pie chart surrounding the UMAP plot. Key for individual nucleosome types and respective ‘groups’ (as defined by hierarchical clustering) are shown on bottom. **B.)** ‘Horizon plots’ for 14 distinct structural classes of footprints, including: chromatosomes (Group I), octameric nucleosomes with focal intra-nucleosomal accessibility (Group II-III), symmetrically breathing nucleosomes (Group IV-V), and shorter footprints consistent with hexasomes, tetramers, and dimers (Group VI-VII). **C.)** ‘Horizon plot’ (left), mean accessibility (middle), and structural visualization (right) of C9 footprints, which are nucleosome core particles accessible at superhelical location [SHL] ±2 (denoted by red asterisk). **D.)** As in (C), but for C5 footprints, which are nucleosome core particles accessible at SHL ±3.5. **E.)** Heatmap of log_2_-transformed odds ratios (ORs) for different nucleosome types across epigenomic domains (H3K4me3-marked promoters, H3K4me1-marked enhancers, H3K27me3-marked Polycomb-repressed chromatin, H3K36me3-marked actively transcribed gene bodies, and H3K9me3-marked constitutive heterochromatin) and repeat elements (mouse major satellite, minor satellite, and telomeric sequences). ORs designated with a grey dot indicate those that are not statistically significant from Fisher’s exact tests (*i.e.* Storey’s *q*-value > 0.1).

Nucleosome types displayed patterns consistent with known structural features of chromatin fibers (summarized in key for **Figure 2A**). Group I types were consistent with chromatosomes (*i.e.* nucleosome core particles associated with linker histone H1), owing to their long footprint sizes and positioning of a short subfootprint in the horizon plot, mirroring the expected additional DNA protection by an H1 molecule; group V, VI, and VII clusters, conversely, were the most accessible footprinted particles, demonstrating patterns consistent with breathable nucleosomes (*i.e.* nucleosomes with a full complement of histone proteins and accessibility at superhelical location [SHL] ±6-7), hexasomes, tetrasomes, and smaller subnucleosomal particles (*e.g.* H2A/H2B dimers; transcription factors), respectively. Finally, we also observed ‘intermediate’ clusters, which demonstrated unique, focal patterns of accessibility within the nucleosome core particle, including at SHL-2 (**Figure 2C**) and at SHL-3.5 (**Figure 2D**). We speculate that these states may reflect specific nucleosome-cofactor interactions (*e.g.* interaction between remodelers^37^ or high-mobility group [HMG] proteins^38^ and SHL-2), or reflect the dynamic assembly/disassembly process between H2A/H2B dimers and the H3/H4 tetramer. The relative abundance of nucleosome types was concordant over multiple independent SAMOSA footprinting experiments (**Extended Data Figure 2D**; median ± MAD Pearson’s *r* = 0.978 ± 0.024), and nucleosome type classification was highly robust to the number of nucleosomes sampled (**Extended Data Figure 2E**). Together these experiments suggest that the majority (86.6% of footprints falling in groups II – VII) of nucleosomes in E14 mESCs display some degree of ‘intranucleosomal’ accessibility, implying pervasive nucleosome distortion epigenome-wide.

We hypothesized that nucleosome types reflect a previously unseen layer of chromatin-based regulation. To test this hypothesis, we quantified differences in the distribution of nucleosome types across the E14 mESC epigenome via a series of Fisher’s Exact Enrichment tests, and visualized resultant data as a heatmap of False Discovery Rate (FDR) corrected effect sizes (**Figure 2E**). Strikingly, we found that distinct nucleosome types were significantly depleted or enriched across a wide range of both mappable epigenomic domains (defined by overlap with published histone modification ChIP-seq data) and unmappable epigenomic regions. In predicted promoter regions (defined by H3K4me3), for instance, we observed significant enrichment for clusters 13 and 14 (indicative of two types of short subnucleosomal protection; cluster 13 odds ratio [C13 O.R.] = 1.670, *q* = 1.16 x 10^-294^; C14 O.R. = 1.856, *q* < 1.79 x 10^-308^). In mouse minor satellite sequence (representing the centromeric region), we observed a significant enrichment for clusters 2 and 7 (indicative of chromatosomes and nucleosomes accessible at SHL3.5; C2 O.R. = 2.468, *q* < 1.79 x 10^-308^; C7 O.R. = 2.559, *q* < 1.79 x 10^-308^). Importantly, these enrichment patterns were quantitatively robust over multiple independent replicate SAMOSA experiments (**Extended Data Figure 2F**), suggesting that these are *bona fide* biological differences in the representation of nucleosome types across the epigenome. These analyses suggest that epigenome-specific processes regulate nucleosome types.

### Single-molecule co-occupancy of structurally distorted nucleosomes

A critical advantage of single-molecule footprinting approaches is their unique ability to measure the molecular co-occurrence of protein-DNA interactions^39,40^. Owing to the multi-kilobase length-scale of SAMOSA footprinting data, IDLI extends such analyses to the co-occupancy of several nucleosomes per fiber. To take advantage of this, we next assessed whether nucleosome types display statistically enriched co-occurrence at single-molecule resolution. We implemented an analysis pipeline (**Extended Data Figure 3A**), wherein we computed the likelihood that trinucleosome stretches contain a greater number of co-occurring nucleosome groups than expected by chance. We visualized test results in two ways: first, as a heatmap of effect sizes (**Extended Data Figure 3B**), and second, as waterfall plots of these effect sizes, ranked and colored by FDR-corrected statistical significance (**Extended Data Figure 3C**). Our analysis revealed striking and significant patterns of co-occupancy at the ‘group’ level, including a marked enrichment of ‘homotypic’ arrangements specifically within active versus repressed epigenomic domains (*e.g.* significant enrichment of VI-VI-VI trinucleosomes in H3K4me1 chromatin; O.R. = 3.189, *q* = 4.48 x 10^-3^). Our analysis also uncovered triplet arrangements that were found enriched on both active and facultatively repressed chromatin, including a symmetric pattern of IV-V-IV (cartoon shown in **Extended Data Figure 3D**). Single-molecule visualization of fibers falling within each of four different chromatin types (**Extended Data Figure 3E**) highlight the flexibility of these analyses, while illustrating trinucleosomal co-occurrence motifs at single-molecule resolution and at any desired sequence genome-wide. Together, these analyses highlight the technical utility of nucleosome classification in uncovering context-specific structural motifs on chromatin fibers. Further, our proof-of-concept reveals specific co-occurrence patterns that are both unique and shared across diverse epigenomic regions.

### The interplay between nucleosome type, transcription factors, and DNA sequence

We next sought to study nucleosome types in the context of TFs and their cognate DNA sequence motifs. We reasoned that the position and identity of nucleosome types in reference to TFs could be analyzed in two complementary ways (**Figure 3A**): first, we could assess the relative abundance of nucleosome types surrounding bound TF motifs compared to unbound controls (*i.e.* a ‘footprint-centric’ view of nucleosome structure analogous to analyses above); second, we could assess the relative abundance and positioning of IDLI footprints / subfootprints centered at motifs themselves (*i.e.* a ‘motif-centric’ view of nucleosome distortion). These analyses relate different structural aspects: footprint-centric views are agnostic to motif position but provide a holistic view of nucleosome structure encompassing motifs (**Figure 3B**); motif-centric views quantify how DNA sequence positions footprints, but only capture the substructure of regions of a footprint overlapping with the motif (**Figure 3C**). Interpreting IDLI footprints in both ways is essential, as how TFs engage their motifs in the context of nucleosomes remains contentious^15,41,42^. Finally, to query whether TFs directly control nucleosome types, we incorporated small-molecule sensitive degron perturbation^43^ within this experimental framework (**Figure 3D**).

**Figure 3:**
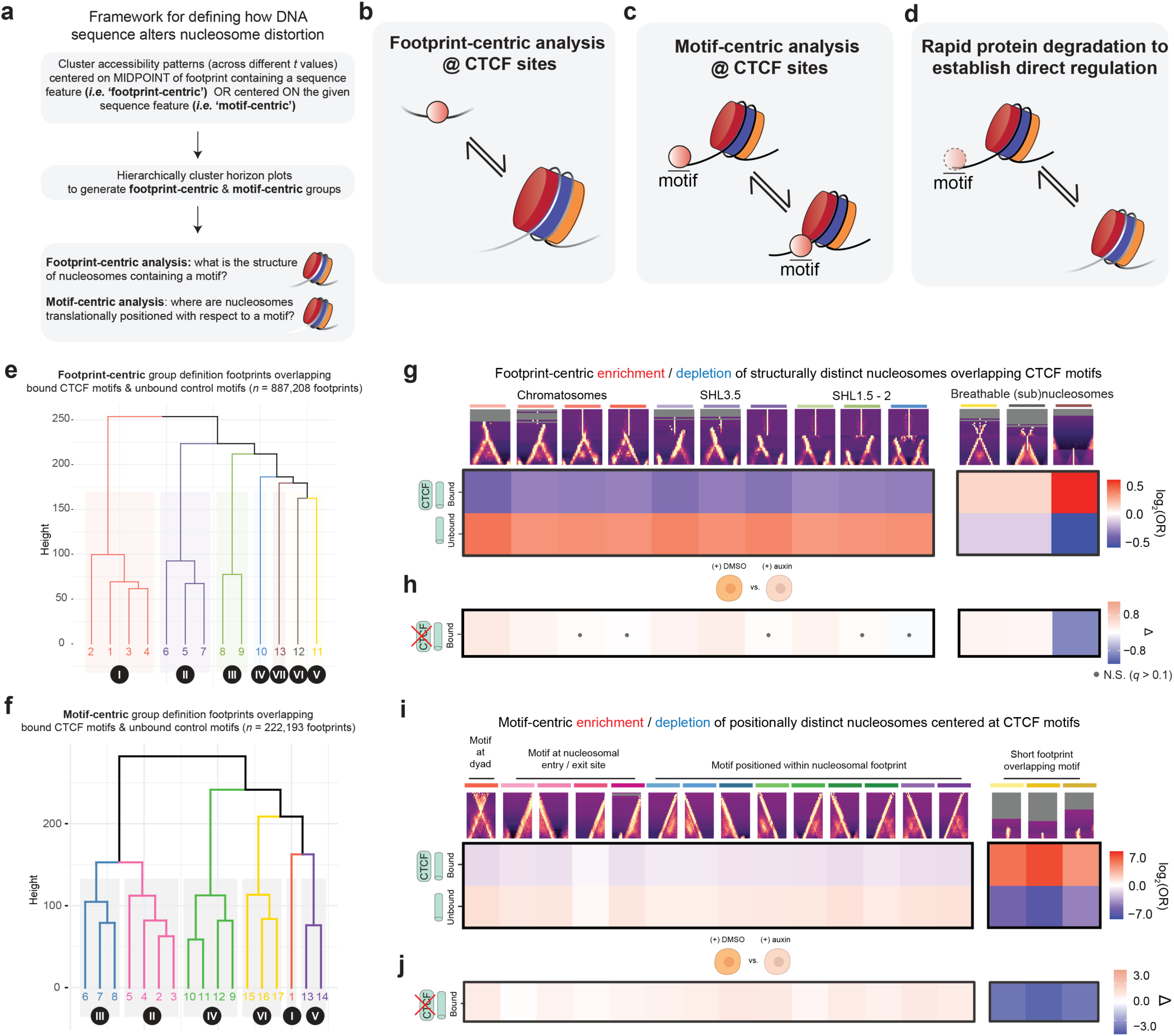
Footprint- and motif-centric analyses of nucleosomes at CTCF motifs in wild-type and CTCF-depleted mESCs. **A.)** Framework for defining nucleosome types and translational positions for footprints at TF-binding motifs of interest. Footprint-centric analyses (*i.e.* clustering accessibility centered on footprint midpoints) quantify distortion patterns for nucleosomes within a 1-kb window centered on a given motif of interest. Motif-centric analyses (*i.e.* clustering accessibility centered on motif sequence) address how nucleosomes are translationally positioned with respect to a given motif. **B-C.)** Footprint-centric and motif-centric analyses were performed in E14 mESC data by comparison of bound vs. randomly sampled motifs for CCTCC-binding factor (CTCF). **D.)** Similar quantification was performed at bound CTCF motifs in mESCs expressing an AID-tagged form of CTCF by comparison of distortion patterns and translational positions before vs. after auxin-mediated CTCF degradation (for 6 hr). **E-F.)** Hierarchical clustering of ‘horizon plot’ data from footprint-centric and motif-centric analyses at bound and unbound CTCF motifs reveals 7 distinct ‘groups’ of nucleosome types and 6 distinct ‘groups’ of translational positions. **G.)** Heatmap of log_2_-transformed odds ratios (ORs) for assessing enrichment/ depletion of different nucleosome types at bound vs. randomly sampled CTCF motifs in E14 mESCs. ORs designated with a grey dot indicate those from Fisher’s exact test that are not statistically significant (*i.e.* Storey *q*-value > 0.1). **H.)** Heatmap of effect sizes (Δ) for nucleosome types at bound CTCF motifs in auxin vs. DMSO-treated CTCF-AID mESCs. Effect sizes designated with a grey dot indicate those from Fisher’s exact test that are not statistically significant (*i.e.* Storey *q*-value > 0.1). **I-J.)** As in **(G-H)**, but for CTCF motif-associated translational positions.

We applied our approach first to the essential TF CCCTC-binding factor (CTCF), which plays diverse roles within the nucleus and has a number of features that make it readily amenable for this analysis: its long residence time^44^, ease-of-footprinting^45^, and well-defined set of E14 mESC binding sites^46^. Importantly, our analysis of CTCF serves as a valuable control to validate our ability to detect different nucleosome configurations (and potentially directly footprint TF binding itself) at sequence-specific recognition motifs. After integrating single-molecule footprinting data from CTCF degron experiments^33,47^ (reproducibility of degradation experiments summarized in **Extended Data Figure 4A-E**), we clustered IDLI footprints using both footprint-centric (nucleosome types) and motif-centric pipelines, resulting in 13 (**Extended Data Figure 4F**; F1-F13) and 17 clusters (**Extended Data Figure 4G**; M1-M17), respectively, which we further grouped through hierarchical clustering (**Figure 3E-F**). Cluster abundances for both footprint-centric and motif-centric CTCF analyses were highly reproducible (**Extended Data Figure 4H-I**).

Footprint-centric clusters qualitatively matched nucleosome types defined in epigenome-wide analyses, demonstrating patterns consistent with chromatosomes, intranucleosomally accessible particles, breathable nucleosomes, subnucleosomes, and short protections (**Extended Data Figure 4F**). We computed the extent to which these nucleosome types differed at ChIP-backed CTCF motifs versus random motif matches unlikely to bind CTCF (**Figure 3G**; reproducibility of enrichment shown in **Extended Data Figure 4J**). We observed significant enrichment of subnucleosome and short footprints at bound CTCF motifs specifically (*e.g.* F13 O.R. = 1.309; *q* < 1.79 x 10^-308^; Fisher’s Exact Test), accompanied by significant depletion for less accessible nucleosome types. Depletion of CTCF protein (**Figure 3H**) resulted in significant loss of short footprints (degron F13 O.R. = 0.681; *q* = 6.93 x 10^-103^; reproducibility of effect size shown in **Extended Data Figure 4K**). These analyses suggest that CTCF protein dictates the composition of structurally distinct nucleosomes (specifically, to favor partially unwrapped states), and that CTCF can generate its own short footprint. Further, acute depletion demonstrates that CTCF directly regulates the abundance of short footprints only, as its loss results in a concomitant gain of all other nucleosome types.

We repeated a similar set of analyses for motif-centric clusters (**Extended Data Figure 4G**). These clusters fell into groups that could be broadly categorized as: i.) nucleosomes of varying breathability with a dyad centered over the CTCF motif (group I); ii.) translationally-positioned nucleosomes with the CTCF motif precisely placed at the nucleosome entry / exit (group II); iii.) nucleosomes with the CTCF motif occurring within the nucleosome core particle (groups III, IV); and iv.) short footprints of varying sizes, precisely positioned over the CTCF motif (group V; possibly reflective of multi-modal binding by the constituent zinc fingers of CTCF^45,48^). Enrichment analysis of these clusters at bound versus control motifs (**Figure 3I**; reproducibility of enrichment shown in **Extended Data Figure 4L**), revealed that group V footprints were significantly enriched at true sites versus control (*e.g.* M15 O.R. = 30.939; *q* < 1.79 x 10^-308^), and degradation experiments (**Figure 3J**; reproducibility of effect size shown in **Extended Data Figure 4M**) confirmed that these short footprints were generated by CTCF itself (degron M15 O.R. = 0.186; *q* < 1.79 x 10^-308^). Together, these analyses establish a technical framework for studying nucleosome heterogeneity at TF binding sites, suggest that there are no specific preferred nucleosome orientations surrounding occupied CTCF motifs, and demonstrate that CTCF binding is the predominant driver of motif-dependent footprint positioning in E14 mESCs.

### SOX2 protein directly regulates nucleosome distortion at cognate binding sites

While strong CTCF motifs are sufficient to outcompete well-positioned nucleosomes in synthetic contexts^49^, how motifs for the pluripotency factor SOX2 contend with nucleosomes remains unclear. Genomic, structural, and other biochemical assays have offered conflicting results on how SOX2 engages chromatin, particularly regarding SOX2 interactions within the nucleosome^50,51^, versus at the nucleosome entry / exit site^49,52^. To provide molecular insight into these possible binding modes for SOX2, we repeated the analyses above at SOX2-OCT4 composite motifs^53^, including data from a SOX2-dTAG mESC line^54^ (degradation characterization and reproducibility in **Extended Data Figure 5A-F**). Footprint-centric analyses revealed 14 nucleosome types (F1-F14), which fell into similar groups as above (**Figure 4A**; **Extended Data Figure 5G**; reproducibility of cluster abundances shown in **Extended Data Figure 5I**). Differential analysis of nucleosome types at bound versus control motifs (**Figure 4C**; reproducibility of enrichment shown in **Extended Data Figure 5K**) revealed a significant enrichment for breathing nucleosomes, subnucleosomes, and short footprints at bound motifs (*e.g.* F13 O.R. = 1.420; *q* = 1.42 x 10^-247^), with depletion of inaccessible nucleosomal species. We speculate that these patterns may reflect the activities of ATP-dependent chromatin remodeling complexes and / or SOX2 altering nucleosome structure at these *cis-*regulatory elements. Like CTCF depletion, acute degradation of SOX2 protein (**Figure 4D**; reproducibility of effect size shown in **Extended Data Figure 5L**) resulted in significant depletion of short footprints (degron F14 O.R. = 0.871, *q* = 1.79 x 10^-3^). Paradoxically, acute depletion of SOX2 is also accompanied by a significant decrease in the proportion of inaccessible nucleosome species (*e.g.* degron F9 O.R. = 0.713, *q* = 1.95 x 10^-5^) in the vicinity of cognate binding motifs. This result reveals an unexpected role for SOX2 protein in binding free DNA to generate short footprints and stabilizing nucleosomal species in proximal regions, possibly by reducing the extent of remodeling at loci with SOX2-OCT4 motifs.

**Figure 4:**
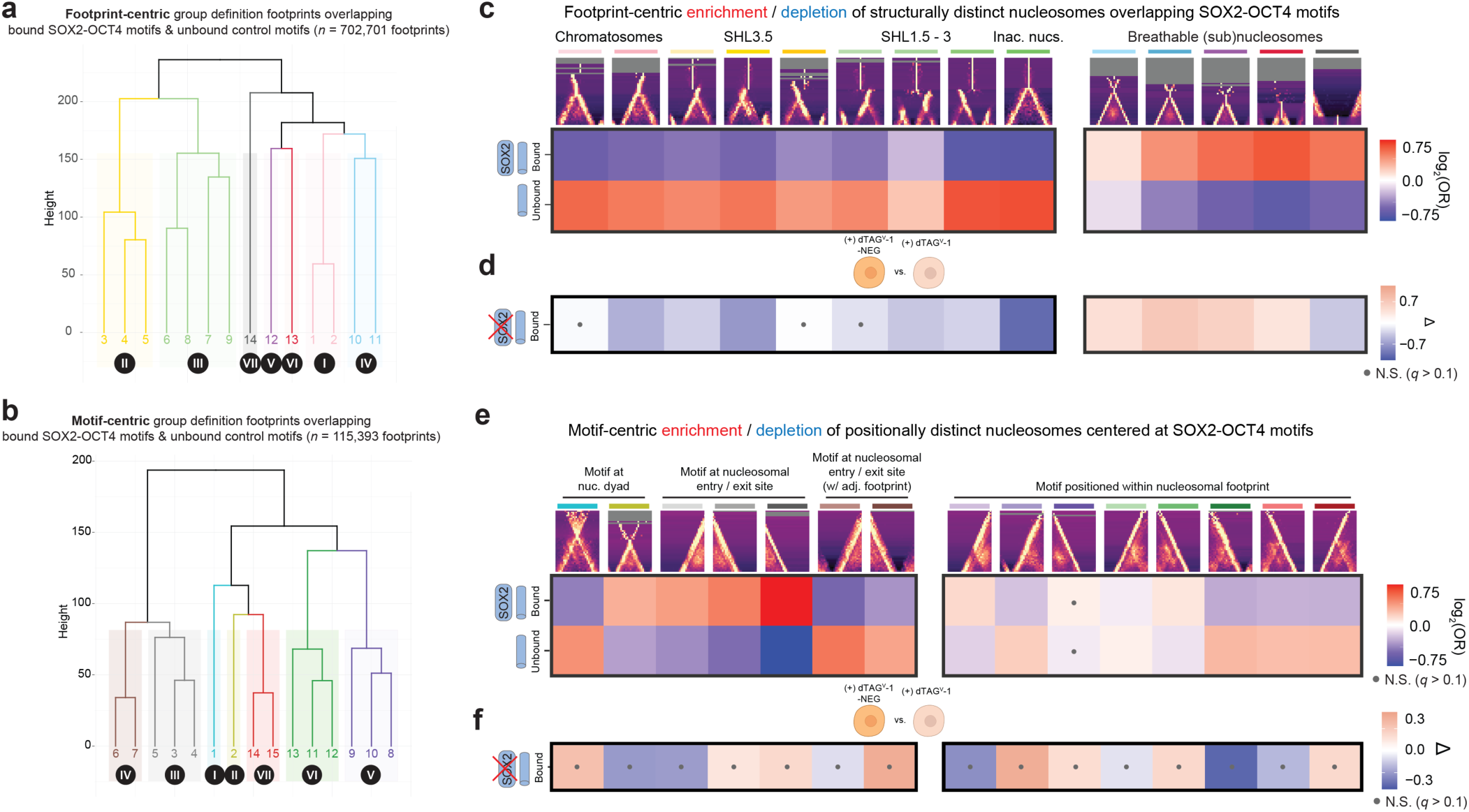
Footprint- and motif-centric analyses of nucleosomes at SOX2-OCT4 motifs in wild-type and SOX2-depleted mESCs. **A-B.)** Hierarchical clustering of ‘horizon plot’ data from footprint-centric and motif-centric analyses at bound and unbound SOX2-OCT4 composite motifs reveals 7 distinct ‘groups’ of nucleosome types and 7 distinct ‘groups’ of translational positions. **C.)** Heatmap of log_2_-transformed odds ratios (ORs) for assessing enrichment / depletion of different nucleosome types at bound vs. randomly sampled SOX2-OCT4 composite motifs in E14 mESCs. ORs designated with a grey dot indicate those from Fisher’s exact test that are not statistically significant (*i.e.* Storey *q*-value > 0.1). **D.)** Heatmap of effect sizes (Δ) for nucleosome types at bound SOX2-OCT4 composite motifs in dTAG^V^-1 vs. dTAG^V^-1-NEG-treated (control) SOX2-dTAG mESCs. Effect sizes designated with a grey dot indicate those from Fisher’s exact test that are not statistically significant (*i.e.* Storey *q*-value > 0.1). **E-F.)** As in **(C-D)**, but for SOX2 motif-associated translational positions.

To further investigate this potential multimodality, we next performed a motif-centric analysis at these sites. Motif-centric clustering yielded 15 different clusters (M1-M15) falling in seven broad groups (**Figure 4B**) with patterns similar to those observed at CTCF sites (**Extended Data Figure 5H**; reproducibility of cluster abundances shown in **Extended Data Figure 5J**). Enrichment analyses at bound motifs compared to unbound controls revealed a striking pattern of translationally positioned nucleosomes, subnucleosomes, and short footprints with respect to bound motifs (**Figure 4E**; reproducibility of enrichment shown in **Extended Data Figure 5M**), and further analysis in the context of protein degradation (**Figure 4F**) revealed no significant effect of the protein itself on these distributions. Importantly, these findings are corroborated by orthogonal studies: enrichment of what appears to be a breathable nucleosome precisely centered over the motif (C2 O.R. = 1.238, *q* = 3.32 x 10^-13^) aligns with recently published genomic mapping studies centered on OCT4, demonstrating that OCT4 can bind an altered nucleosome core particle directly^55^. Moreover, the most abundant enriched states (C3-C5) harbor a motif precisely placed at the nucleosome entry / exit site, consistent with work demonstrating this preference^52^. Finally, the lack of significant effect of SOX2 degradation on these trends implies that additional regulatory factors must be required to reposition nucleosomes and enable SOX2 binding, consistent with recent genomic studies perturbing SWI/SNF remodelers^56,57^. Taken together, these analyses suggest multiple modes of interaction between SOX2 and nucleosomes at bound motifs; motif-encoded interactions with breathable (or noncanonical) nucleosome core particles and nucleosome edges occur concurrently with motif-independent interactions with stable nucleosome core particles.

### Widespread and regulated nucleosome distortion *in vivo*

We next sought to determine if the landscape of regulated nucleosome types exists beyond cultured mESCs, in a tissue *in vivo*. We generated reproducible single-molecule accessibility data in purified mouse hepatocytes for *n* = 7 mice, including a published genetic model^58^ heterozygous for a *Foxa2* allele lacking a critical nucleosome-interacting domain (*i.e.* ‘FOXA2-ΔHx’; characterization and reproducibility of heterozygous deletion in **Extended Data Figure 6A-E**). We then performed footprint-centric IDLI analyses (motif-centric tests all insignificant and not shown) on footprints drawn from regions surrounding ChIP-backed FOXA2 binding sites^59^ and matched negative controls, to assess whether nucleosome types exist in primary cells.

Footprint-centric analyses yielded 15 nucleosome types (F1-F15) / seven groups, corresponding qualitatively to structurally distinct (sub)nucleosomal protections seen in all mESC analyses (**Figure 5A**; **Extended Data Figure 6F**). This suggests that widespread nucleosome distortion is a general property of mammalian chromatin, even within a terminally differentiated primary cell type. Computing the relative enrichment of these clusters at ChIP-backed FOXA2 binding sites compared to random motif matches (**Figure 5B**; reproducibility of enrichment shown in **Extended Data Figure 6G**), we observed a similar TF-dependent pattern to that seen for SOX2 in mESCs, with breathable (sub)nucleosomes and short footprints significantly enriched (*e.g.* F12 O.R. = 2.284, *q* = 2.16 x 10^-68^), and inaccessible nucleosomes significantly depleted at *bona fide* binding sites. Differential nucleosome type analysis in FOXA2-ΔHx heterozygous hepatocytes versus wildtype littermate controls revealed significant depletion of two nucleosome types: F7 (predicted accessibility at SHL-3.5; mutant O.R. = 0.624, *q* = 6.45 x 10^-11^) and F11 (one of 5 chromatosome-like types; mutant O.R. = 0.820, *q* = 8.02 x 10^-4^), providing further evidence that the FOXA2 Hx motif controls nucleosome structure (reproducibility of effect size shown in **Extended Data Figure 6H**. Taken together, these analyses demonstrate the *in vivo* relevance of nucleosome distortion at cell-type-specific regulatory elements, and illustrate multimodal regulation of nucleosome structure by a histone-binding domain of FOXA2.

**Figure 5:**
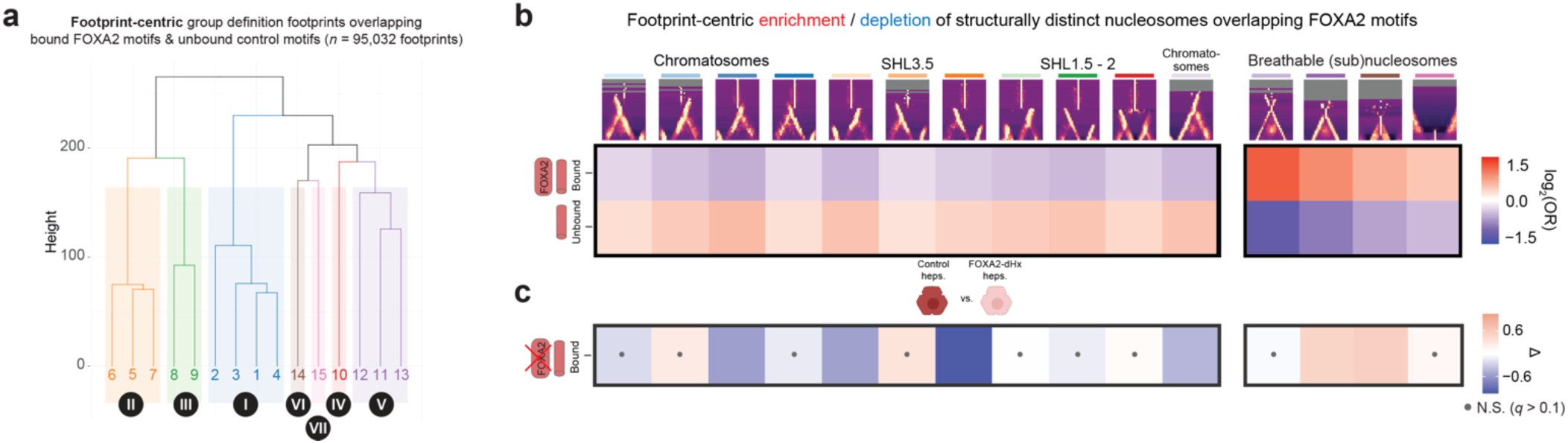
Footprint-centric analyses of nucleosomes at FOXA2 motifs in wild-type primary hepatocytes and hepatocytes derived from heterozygous FOXA2-ΔHx mice. **A.)** Hierarchical clustering of ‘horizon plot’ data from footprint-centric analyses at bound and unbound FOXA2 motifs reveals 7 distinct ‘groups’ of nucleosome types. **B.)** Heatmap of log_2_-transformed odds ratios (ORs) for assessing enrichment / depletion of different nucleosome types at bound vs. randomly sampled FOXA2 motifs in C57BL/6J hepatocytes. ORs designated with a grey dot indicate those from Fisher’s exact test that are not statistically significant (*i.e.* Storey *q*-value > 0.1). **C.)** Heatmap of effect sizes (Δ) for nucleosome types at bound FOXA2 motifs in heterozygous FOXA2-ΔHx hepatocytes vs. wild-type hepatocytes (derived from age-matched littermate controls). Effect sizes designated with a grey dot indicate those from Fisher’s exact test that are not statistically significant (*i.e.* Storey *q*-value > 0.1).

## DISCUSSION

We report the discovery of pervasive, patterned, and programmed nucleosome distortion across mammalian epigenomes (**Figure 6**). Using a newly developed analytical pipeline we term “IDLI” we reveal that the majority of nucleosomes across diverse epigenomic states exist in a partially accessible state (**Figure 6A**), ranging from focal accessibility at specific sites along the nucleosomal wrap, to symmetrically unwound / breathable nucleosomes and subnucleosomes. Our pipeline also classifies short DNA protections – given the raw abundance of these protections, these are most likely owed to the histone H2A/H2B dimer, though our analyses at TF binding sites show a quantifiable (and expected) contribution of TFs to the overall number of short protections.

**Figure 6:**
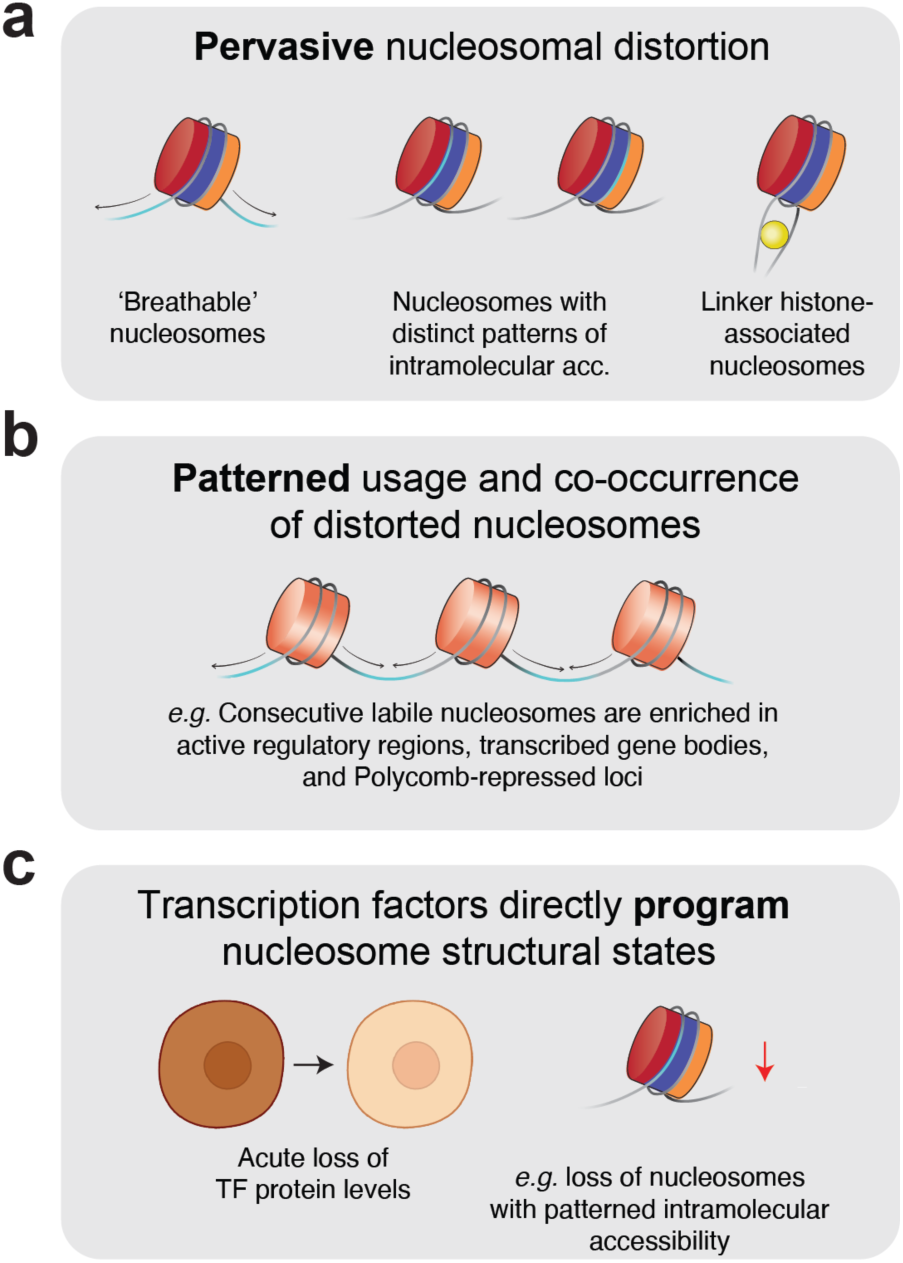
IDLI reveals pervasive, patterned, and programmed nucleosome distortion within mammalian chromatin. **A.)** Analyses of intranucleosomal accessibility from non-destructive footprinting data indicate that most nucleosomes surveyed are not entirely refractory to EcoGII methylation; instead, nucleosomes are often symmetrically ‘unwound’ or highly ‘breathable’, they can exhibit focal accessibility at specific superhelical locations, or they can exist as chromatosomes. **B.)** We find that patterns of nucleosomal distortion are enriched / depleted to different extents across epigenomic domains and repeat elements (not shown here). Moreover, we demonstrated that consecutive homotypic ‘triplets’ of nucleosomes are often statistically enriched on single fibers (*e.g.* ‘breathing’ nucleosomes tend to co-occur on individual chromatin fibers in active and facultatively repressed chromatin). **C.)** Across multiple cell types and genetic perturbations, we uncover that TFs are directly responsible for changes in the abundance of different nucleosome types at loci that harbor their cognate binding motifs. In the context of acute protein depletion (*i.e.* CTCF and SOX2-degron lines), these observations strongly suggest a direct role in TF-mediated programming of nucleosomal states.

IDLI analyses capture regulated patterns of nucleosome distortion genome-wide (**Figure 6B**). Our proof-of-concept study demonstrates that nucleosome types—which we interpret as structurally distinct nucleosomes—are specifically enriched and depleted across epigenomic domains; further, our first-in-class single-fiber co-occupancy analyses demonstrate the existence of statistically-significant co-occurrence of structurally distinct nucleosomes on individual chromatin fibers. These analyses are uniquely enabled by the IDLI pipeline, which operates on long-read high-resolution footprinting data. Future work must assess the significance of these co-occurrence patterns at even higher resolution (*e.g.* at actively transcribed promoters, enhancers, or gene bodies, or at individual TF binding sites, which we have preliminarily computed for CTCF and SOX2 in mESCs to demonstrate feasibility; **Extended Data Figure 7**). Given the short residence times of most TFs^60^, however, we stress that a comprehensive TF-specific co-accessibility analysis will require SAMOSA data with increased genomic coverage specifically at target TF binding sites. In the future, this may be achieved via additional technology improvements (*e.g.* higher-throughput multiplex target capture of TF binding sites^61^, analogous to multiplex bisulfite sequencing^40,62^).

Finally, we show that the TFs CTCF, SOX2, and FOXA2 directly program nucleosome distortion at cognate binding sites (**Figure 6C**). Leveraging small molecule sensitive degrons (CTCF; SOX2) and a mutant *Foxa2* allele defective for histone binding, we provide compelling genetic evidence that TF dosage directly influences the structure and positioning of nucleosomes at cognate motifs in cell culture and *in vivo*. Our results for SOX2 and FOXA2 imply multiple modes by which TFs interact with DNA motifs and / or nucleosomes. For both TFs, depletion (or mutation) paradoxically results in a loss of less-accessible nucleosome types, signaling a direct role for these factors in stabilizing nucleosomes. In the case of SOX2 specifically, this phenotype appears to be motif-independent; motif-centric analyses, conversely, demonstrate a preferred positioning state of nucleosomes with the motif oriented at the nucleosome entry / exit site. Our results thus harmonize seemingly conflicting results in the field regarding SOX2 binding. First, binding and nucleosome positioning by SOX2 is clearly motif-encoded, and SOX2 is not sufficient to dictate nucleosome positioning at binding sites; these results are consistent with studies demonstrating the importance of ATP-dependent chromatin remodeling to SOX2 motif accessibility^56,57,63^, as well as studies demonstrating the inability of SOX2 to overcome nucleosomal occlusion^49,52^. Second, SOX2 itself can stabilize nucleosomes in the vicinity of SOX2-OCT4 motifs; we speculate that this ‘on-nucleosome’ regulatory mode may relate to observed interactions between SOX2 and the nucleosome dyad^50^, and further note structural evidence that SOX2 and other HMG-box proteins can interact with the nucleosome directly (*e.g.* at SHL2^38,51^, though we acknowledge no specificity for SHL-2 in our data). Finally, these properties appear to be shared at bound FOXA2 motifs in primary mouse hepatocytes. Compared to unbound control motifs, FOXA2-regulated sites are specifically enriched for accessible (sub)nucleosomes and short protections, supporting previous models for a motif-encoded grammar for these interactions^64^. Altering the dosage of intact FOXA2 protein by replacing one allele with FOXA2-ΔHx has a significant impact, however, on the observed distribution of nucleosome types at FOXA2 binding sites, resulting in loss of nucleosomes with accessibility at SHL-3.5, and chromatosomes; these findings are consistent with the canonical definitions of TF pioneering^65^, in which the nucleosome interaction itself instructs cell fate^58^.

Looking ahead, we anticipate the broad applicability of the IDLI pipeline to a wide variety of problems in chromatin biology. ATP-dependent chromatin remodeling^29,30^, nucleosome assembly, transcription^66^, replication^31^, and all other nuclear processes regularly generate structurally distorted nucleosomes, but prevailing approaches to study this distortion rely on *in vitro* biochemical and biophysical assays. Moving forward, we envision a routine experimental framework that complements highly resolved biochemistry with near-nucleotide-precise snapshots of structurally distorted states *in vivo*. Combining bottom-up biochemistry, top-down structural classification presented here, and complementary advances in *in situ* chromatin imaging^67^ will ultimately enable deeper quantitative insight into the *in vivo* biochemistry of essential nuclear regulatory processes.

## EXTENDED DATA FIGURES

**Extended Data Figure 1:**
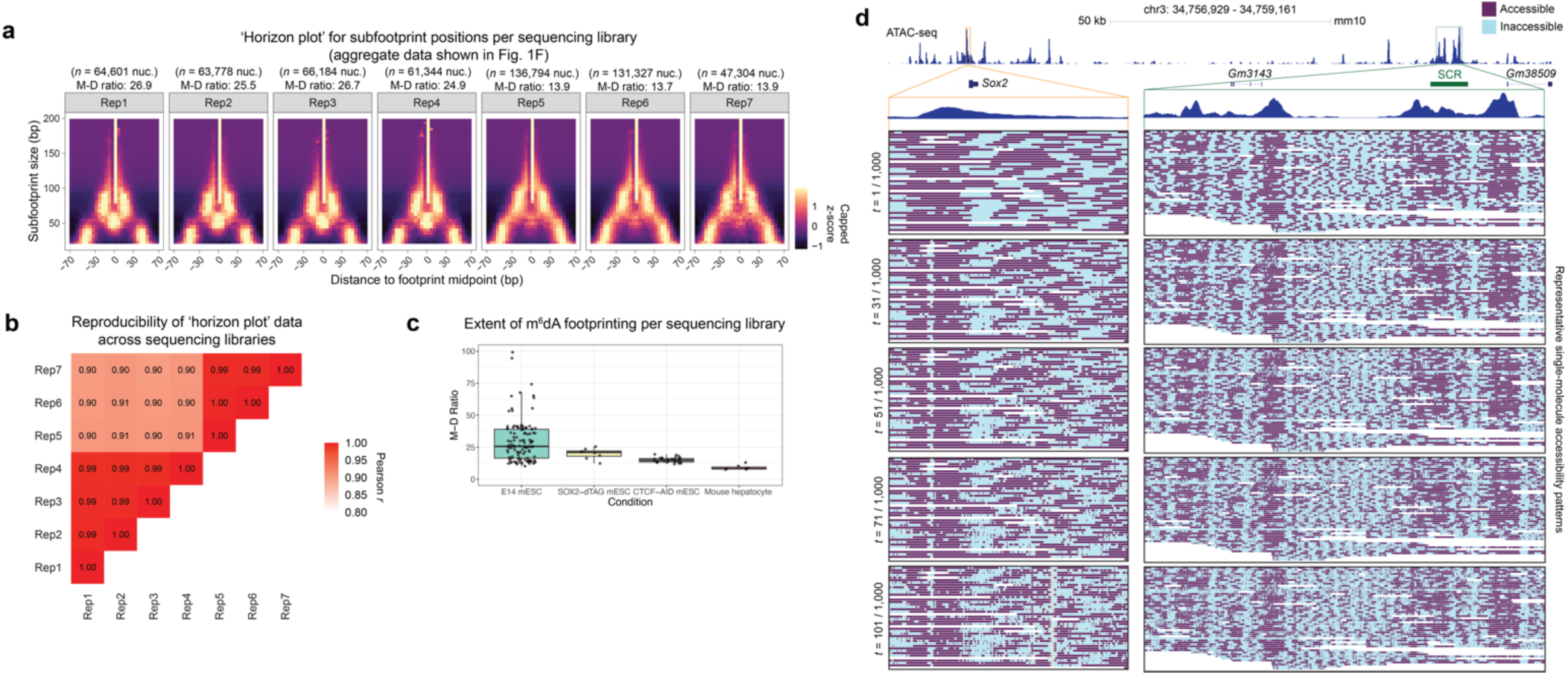
Reproducibility of subfootprint detection with IDLI framework, related to Figure 1. **A.)** ‘Horizon plots’ for enrichment patterns of subfootprints of different sizes at varying positions with respect to footprint midpoints, plotted separately for nucleosomal footprints derived from individual biological replicates (for aggregate data shown in **Figure 1F**). Subfootprints are reliably detected across sequencing libraries with different extents of methyltransferase footprinting, as indicated by mononucleosome-dinucleosome ratio (MDR; value shown above each plot). **B.)** Pearson *r* values from comparison of ‘horizon plot’ data across sequencing libraries (from data shown in **Extended Data Figure 1A**). Samples with more similar MDRs exhibited quantitively higher levels of correlation in reproducibility of ‘horizon plot’ data (*e.g.* Rep1-4 vs. Rep5-7). **C.)** Box-and-whisker plots of MDR values for SAMOSA data included in this study, wherein individual points represent data from single biological replicates. Data are stratified based on cell type and genotype. All libraries from mESCs included in this study are well-footprinted based on previously-defined criteria (*i.e.* MDR > 10). Primary hepatocyte samples (wild-type and FOXA2-ΔHx) have slightly lower MDR values (MDR = 7.53 – 13.06), which we ascribe to slightly reduced efficiency of EcoGII footprinting in primary cells (compared to cell culture conditions) consequent to global differences in chromatin accessibility in pluripotent cells vs. terminally-differentiated cell types. **D.)** Single-molecule accessibility patterns (EcoGII-accessible and -inaccessible DNA in purple and teal, respectively) at *Sox2* promoter (left; depicted in orange) and *Sox2* Control Region [SCR] (right; depicted in green). Data from representative identical single EcoGII-footprinted fibers are shown for the full range of transition probabilities used in IDLI (*t* = 1, 31, 51, 71, and 101 / 1000). Bulk-average chromatin accessibility data (from ATAC-seq) in mESCs are also shown across the entire locus (top) and at specific regulatory loci (bottom).

**Extended Data Figure 2:**
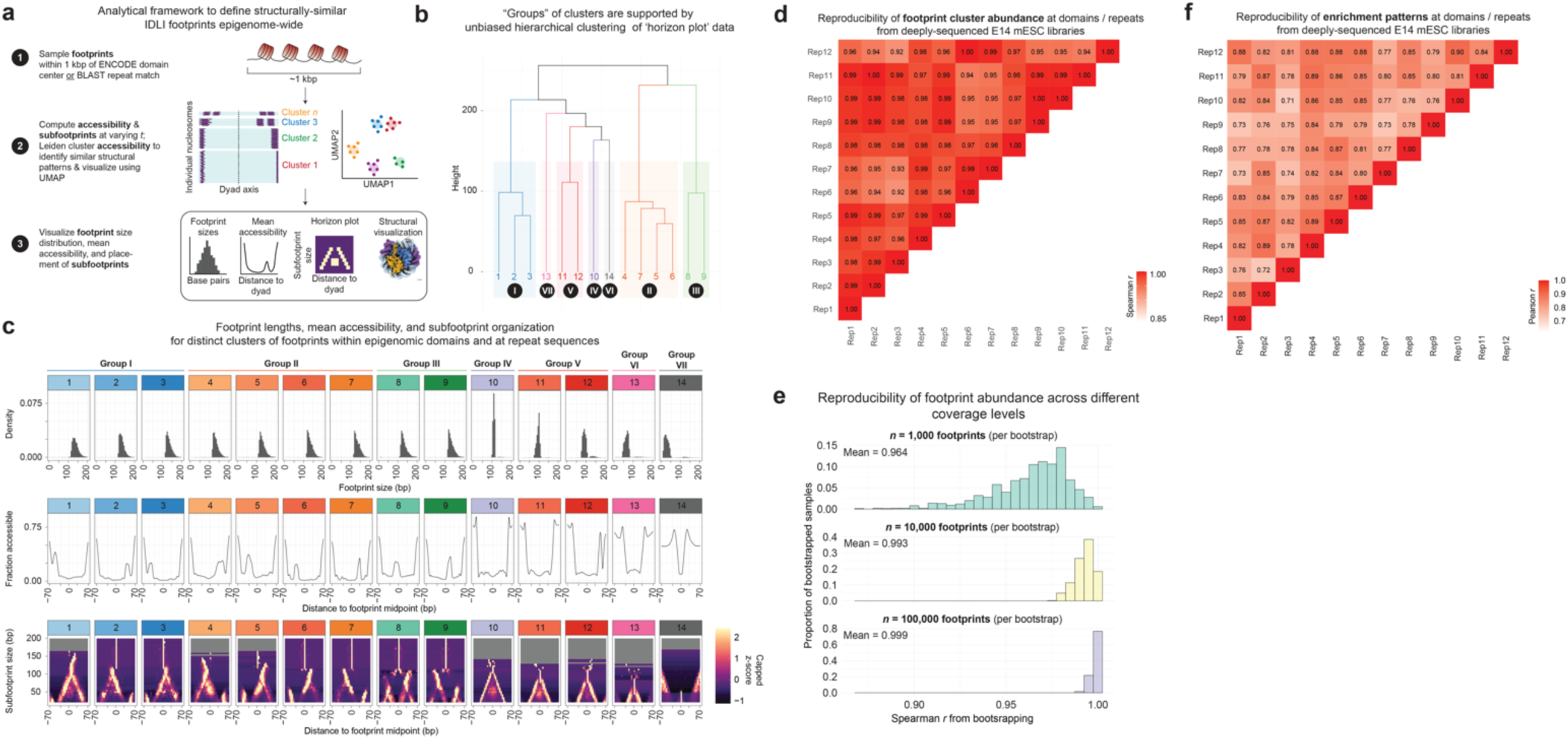
Clustering analysis for defining epigenomic domain- and repeat element-associated nucleosomal distortion patterns, related to Figure 2. **A.)** Framework for clustering and visualization of footprints with shared structural features (*i.e.* patterns of intranucleosomal accessibility) within epigenomic domains and at repeat sequences. Footprints are sampled from within ±500 bp window centered on either the midpoint of ENCODE-defined domains or from the top-scoring repeat for sequencing reads that contain one or more BLAST matches. Single-footprint accessibility patterns across *t* = 31, 51, and 71 / 1,000 are subject to Leiden clustering, and resultant data are shown as a UMAP visualization. To ascribe biological function to footprints with different Leiden assignments, we visualized: (1) footprint size distributions, (2) mean accessibility profiles with respect to footprint midpoints, (3) ‘horizon plots’ for detecting subfootprint placement, and (4) structural visualization of mean accessibility overlaid on the PDB structure for an interphase nucleosome (7KBD). **B.)** Hierarchical clustering of ‘horizon plot’ data from 14 Leiden-defined clusters of domain and repeat-associated footprints into 7 distinct groups with shared intra-nucleosomal accessibility patterns. **C.)** Analysis of individual Leiden-defined clusters of nucleosomal distortion patterns at epigenomic domains and repeat sequences. **Top:** Histograms of footprint sizes on a per-cluster basis. Middle: Mean accessibility traces centered on footprint midpoints on a per-cluster basis. **Bottom:** ‘Horizon plots’ for subfootprint sizes versus distance from footprint midpoints to assess subfootprint enrichment at particular locations on a per-cluster basis. z-scores are capped such that the maximum shown in the heatmap are the 95^th^ percentile values within ±70 bp of the respective footprint midpoints. **D.)** Spearman *r* values from comparison of proportions of nucleosome types at domains and repeats across biological replicates (from *n* = 12 selected, deeply-sequenced mESC samples surveyed for both domain- and repeat-associated footprints). **E.)** Bootstrapping analysis to determine number of footprints required for reproducible cluster assignments. From *n* = 795,765 domain- and repeat-associated footprints, we bootstrapped *n* = 1,000, 10,000, or 100,000 footprints per trial. Shown are the distribution of resultant Spearman *r* values (from a total of 10,000 trials per condition). These data suggest that our Leiden assignments for clusters of nucleosomal distortion patterns are reproducible and robust at the number of nucleosomes we have chosen for analysis. **F.)** Pearson *r* values from comparison of nucleosome type enrichment patterns across domains and repeats (aggregate data shown in **Figure 2E**) across biological replicates (from *n* = 12 selected, deeply-sequenced mESC samples surveyed for both domain- and repeat-associated footprints).

**Extended Data Figure 3:**
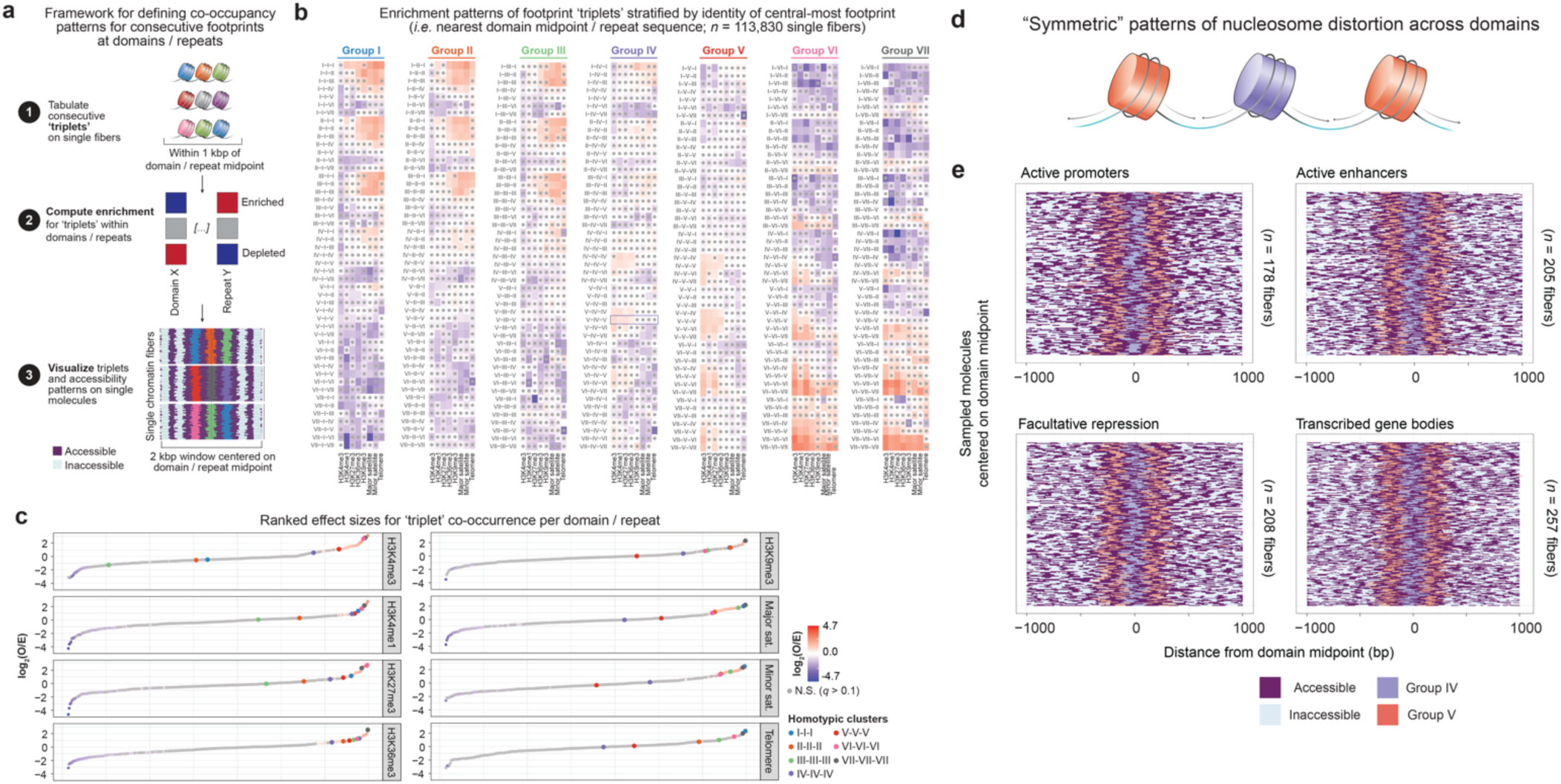
Co-occupancy analysis reveals enrichment of homotypic clusters of footprints with shared nucleosomal distortion patterns on individual fibers. **A.)** Domain- and repeat-associated footprints were analyzed for co-occupancy patterns on single chromatin fibers. Instances of three consecutive footprints on individual sequencing reads nearest domain or repeat midpoints were tabulated based on their associated group assignments (as defined by hierarchical clustering in **Extended Data Figure 2B**). We required that at least three Leiden-classified footprints occurred within ±500 bp of the domain or repeat midpoint of interest, excluding fibers with footprints > 200 nt in length and/or with footprints from the least abundant (bottom 10%) of Leiden-defined clusters. **B.)** Heatmap of log_2_-transformed O / E ratios computed for each possible ‘triplet’ on a per-domain or per-repeat basis from a total of *n* = 113,830 single fibers. Data are stratified based on the group identity of the central-most footprint (*i.e.* footprint nearest the domain / repeat midpoint of interest). Effect sizes designated with a grey dot indicate those from Fisher’s exact test that are not statistically significant (*i.e.* Storey *q*-value > 0.1). **C.)** Waterfall plots of ranked effect sizes stratified by locus indicates enrichment of homotypic clusters of footprints with shared nucleosomal distortion patterns. As in (B), grey dots used to indicate effect sizes for ‘triplets’ with *q* > 0.1. Red and blue dots are used to indicate ‘triplets’ that are enriched and depleted within a given locus, respectively. ‘Triplets’ that comprise homotypic clusters are highlighted by larger colored dots (see legend on bottom right). **D.)** Schematic for ‘triplets’ of symmetric pattern of ‘breathable’ nucleosomes (IV-V-IV), which are statistically enriched in active and facultatively repressed chromatin. **E.)** Single-molecule visualization of chromatin fibers with IV-V-IV ‘triplets’, wherein each row represents a unique sequenced fiber (total number of molecules indicated on right side of each plot). Accessibility patterns across a 2-kb window centered on domain midpoints of interest are also depicted (purple for EcoGII-accessible and teal for EcoGII-inaccessible DNA, respectively). Footprints from different groups are colored as described in associated legend; relative widths depict the number of nucleotides protected per footprint. Only data from epigenomic domains that are significantly enriched for the given triplet (IV-V-IV) are shown.

**Extended Data Figure 4:**
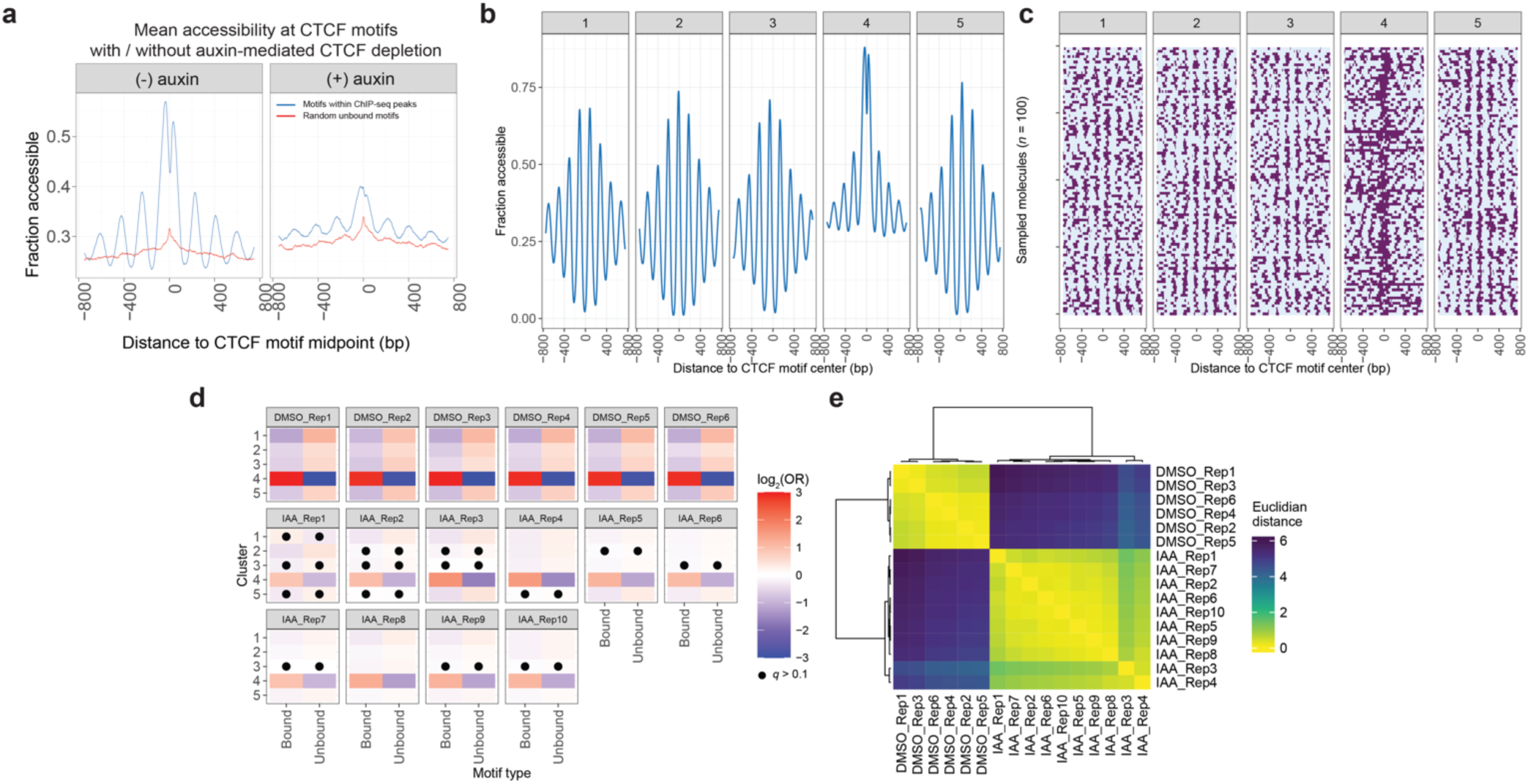
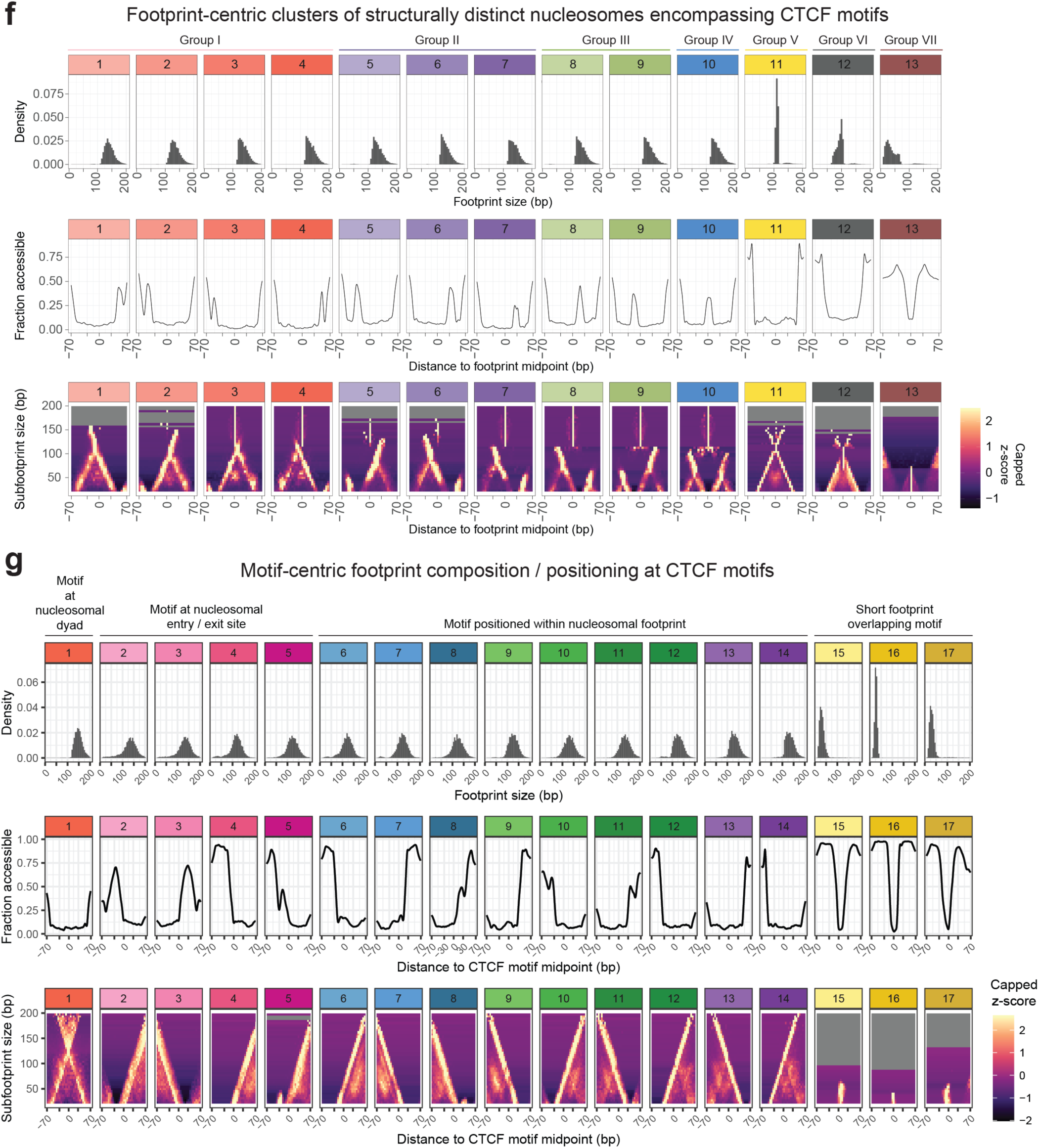
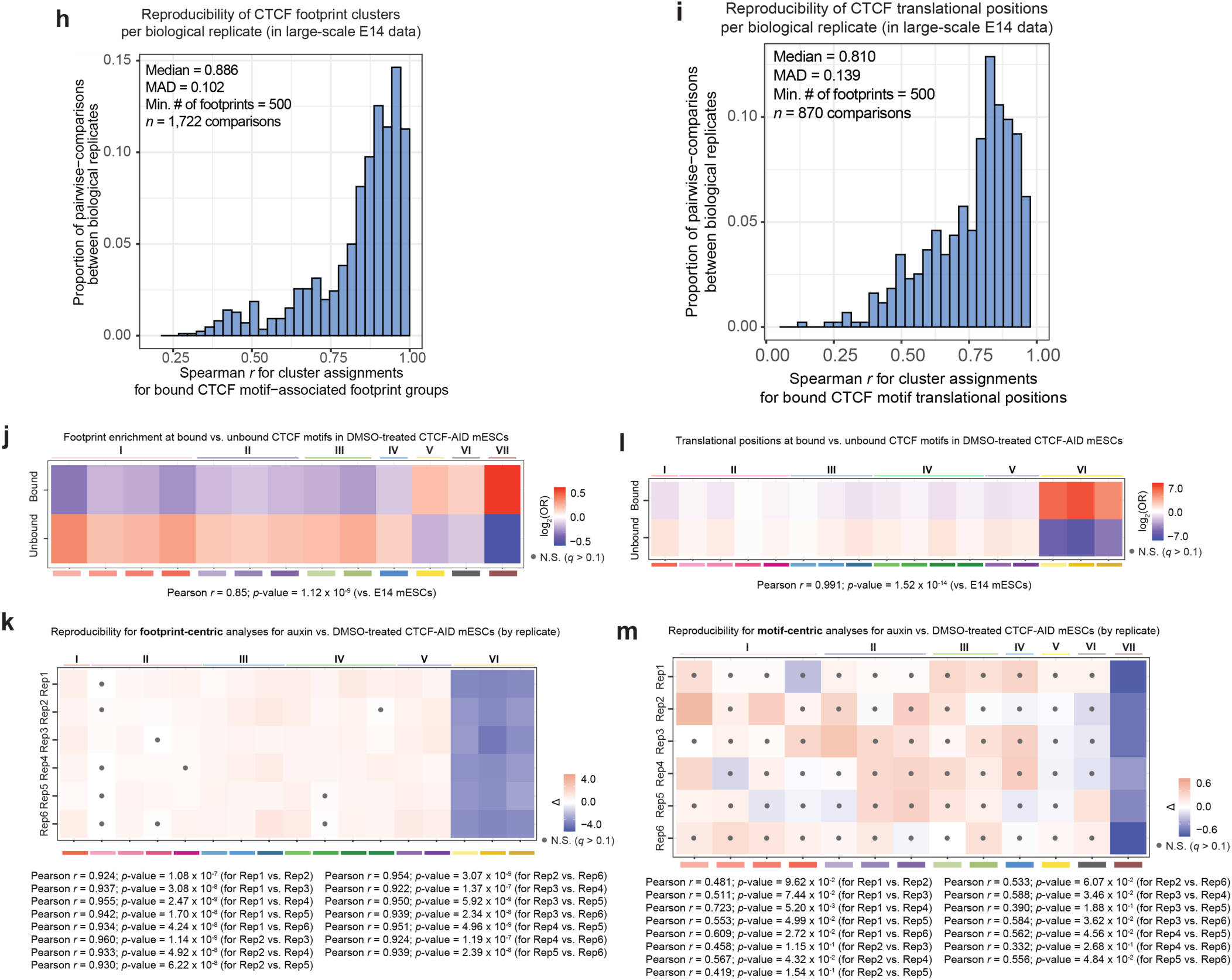
Validation and reproducibility of CTCF-associated distortion patterns and translational positions, related to Figure 3. **A.)** Fraction accessible across ±750 bp window centered on bound (blue) and random (red) CTCF motifs. Comparison of data from DMSO- and auxin-treated CTCF-AID mESCs indicates that acute depletion of CTCF protein is associated with a loss in protection directly at bound CTCF motifs, a decrease in mean accessibility at bound CTCF loci, and an associated loss in phased nucleosomes flanking the nucleosome-free region. **B.)** Line plot representation of mean fraction accessible for Leiden-defined clusters at CTCF motifs in CTCF-AID mESCs. **C.)** Sampled single molecules underlying the averages shown as lineplots for each cluster. Each line represents 1,500 nucleotides extracted from an individual molecule, centered at the CTCF motif. **D.)** Heatmap representation of enrichment (red) or depletion (blue) of clusters comparing fibers at bound vs. unbound CTCF motifs. Data are shown for DMSO- and auxin-treated (designated as ‘IAA) CTCF-AID mESC samples, stratified by biological replicate. Black dots mark Fisher’s exact tests where *q* > 0.1 (not significant). **E.)** Heatmap of Euclidian distances computed from enrichment data in (D); replicates cluster by drug treatment, suggesting that loss of CTCF protein is associated with a reproducible change in accessibility states at CTCF motif loci. **F.)** Analysis of individual Leiden-defined clusters of nucleosome types at CTCF motifs. **Top:** Histograms of footprint sizes on a per-cluster basis. **Middle:** Mean accessibility traces centered on footprint midpoints on a per-cluster basis. Bottom: ‘Horizon plots’ for subfootprint sizes versus distance from footprint midpoints to assess subfootprint enrichment at particular locations on a per-cluster basis. z-scores are capped such that the maximum shown in the heatmap are the 95^th^ percentile values within ±70 bp of the respective footprint midpoints. **G.)** As in (F), but for CTCF-associated translational positions. Accordingly, mean accessibility and ‘horizon plots’ are instead plotted relative to motif positions. **H.)** Distribution of Spearman’s *r* values for pairwise comparisons of cluster abundances for CTCF motif-associated distortion patterns in large-scale E14 mESC dataset. A minimum of *n* = 500 footprints was required in order to exclude lowly-sequenced libraries from this correlation analysis. **I.)** As in (H), but for CTCF-associated translational positions. J.) Heatmap of log_2_-transformed odds ratios (ORs) for assessing enrichment / depletion of different nucleosome types at bound vs. randomly sampled CTCF motifs in DMSO-treated CTCF-AID mESCs. ORs designated with a grey dot indicate those from Fisher’s exact test that are not statistically significant (*i.e.* Storey *q*-value > 0.1). Correlation is computed relative to E14 mESC data (data from **Figure 3G**). **K.)** Heatmap of effect sizes (Δ) for nucleosome types at bound CTCF motifs in auxin vs. DMSO-treated CTCF-AID mESCs, stratified by biological replicate. Effect sizes designated with a grey dot indicate those from Fisher’s exact test that are not statistically significant (*i.e.* Storey *q*-value > 0.1). **L-M.)** As in **(J-K)**, but for CTCF-associated translational positions.

**Extended Data Figure 5:**
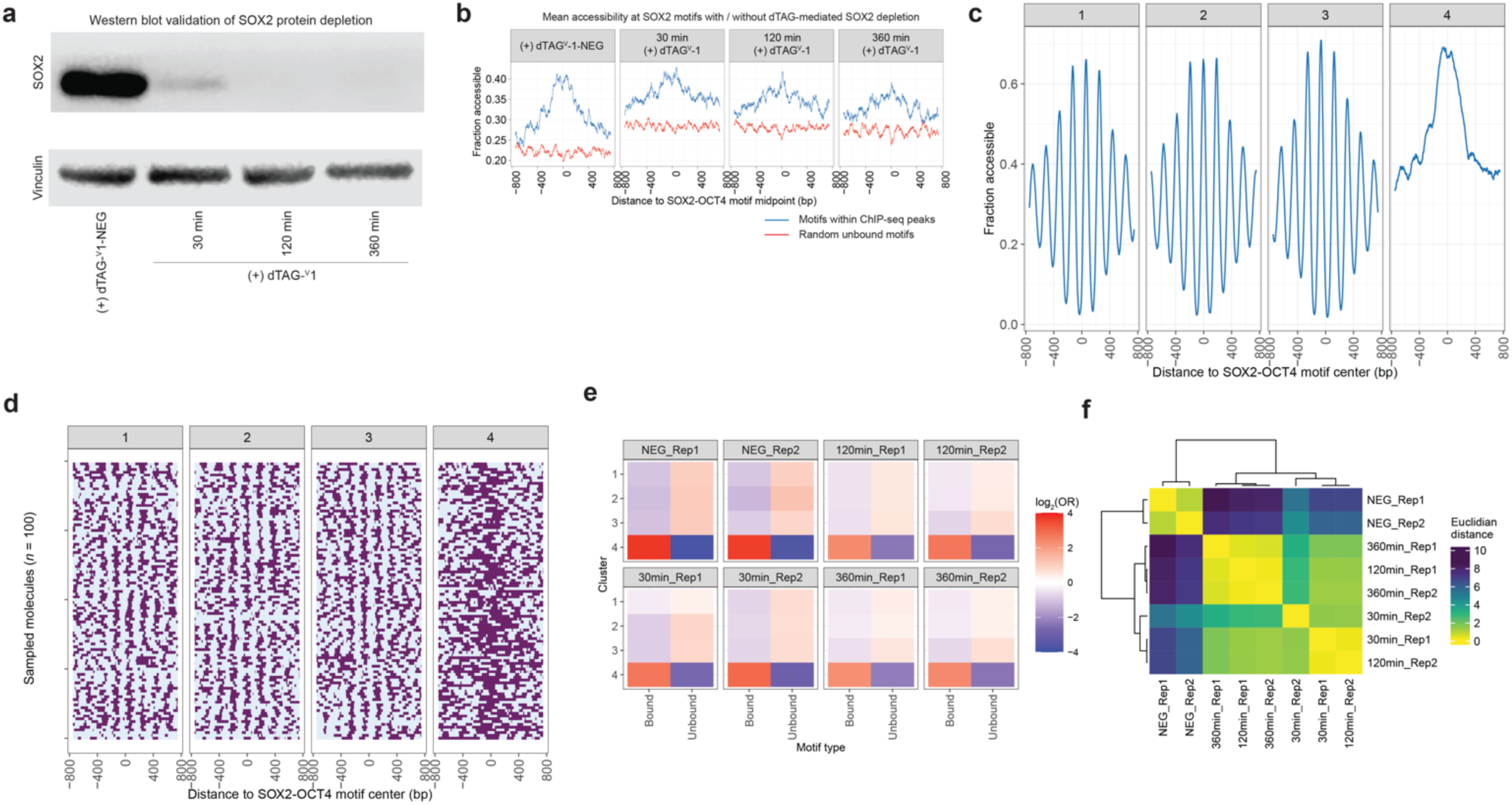
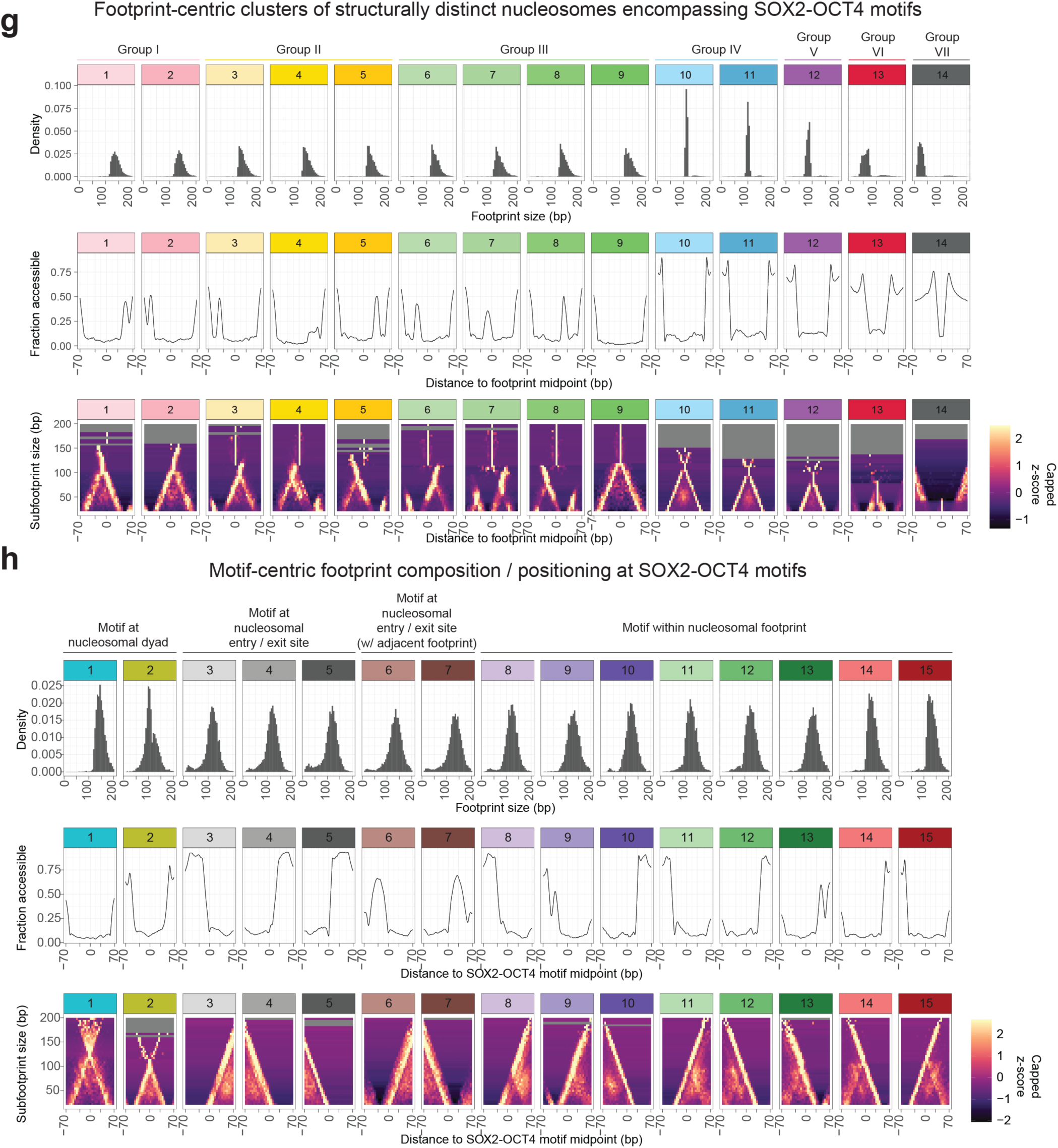
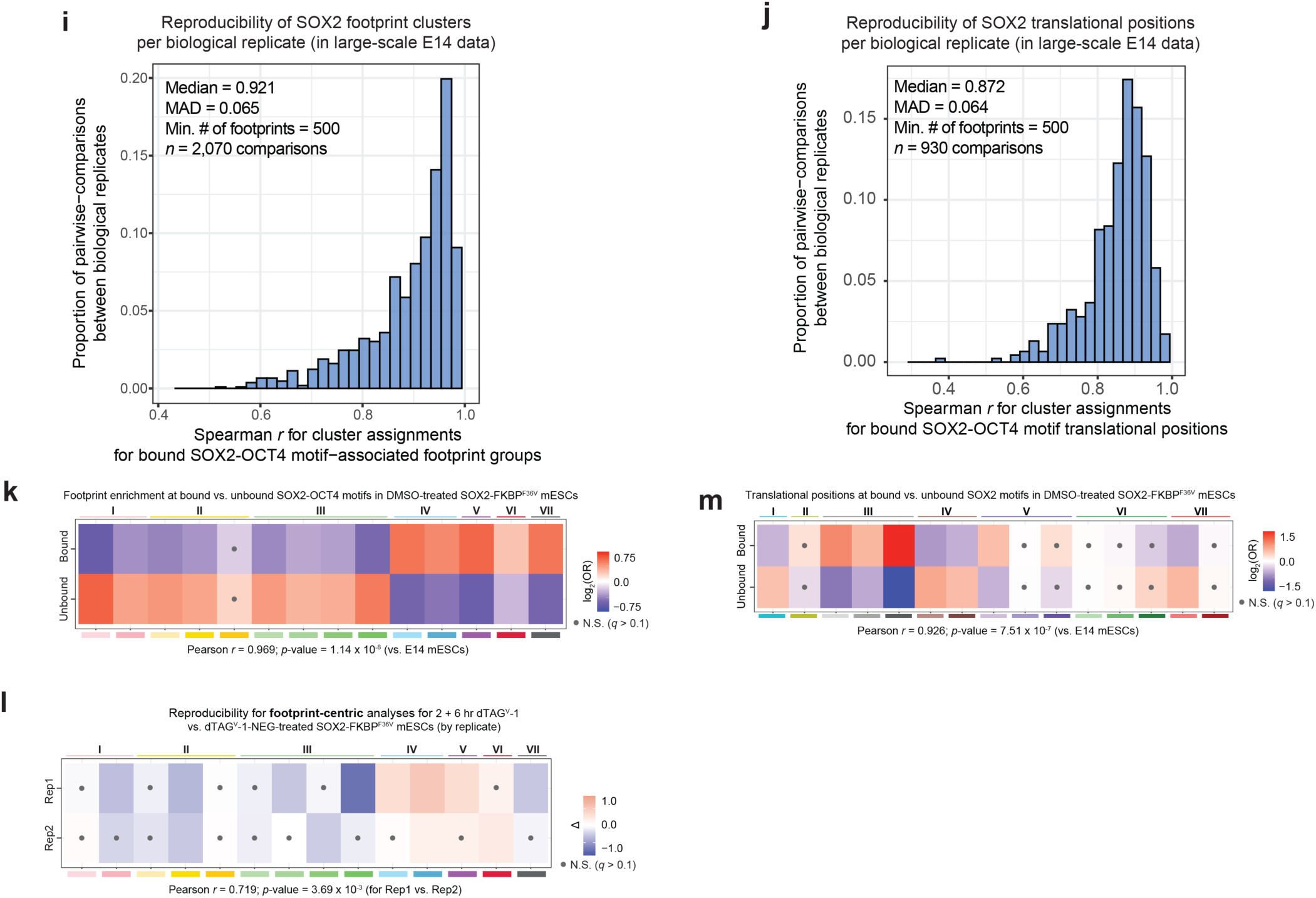
Validation and reproducibility of SOX2-associated distortion patterns and translational positions, related to Figure 4. **A.)** Western blot validation of dTAG-mediated degradation of SOX2 protein indicates near-complete degradation within 30 min of dTAG-^V^1 treatment and undetectable levels of SOX2 by 2 hr and 6 hr. **B.)** Fraction accessible across ±750 bp window centered on bound (blue) and random (red) SOX2-OCT4 composite motifs. Comparison of data from dTAG^V-^1-NEG- and dTAG^V-^1-treated SOX2-dTAG mESCs indicates that acute depletion of SOX2 protein is associated with a slight decrease in chromatin accessibility at SOX2-OCT4 motif loci that fall within ChIP-nexus peaks. **C.)** Line plot representation of mean fraction accessible for Leiden-defined clusters at SOX2-OCT4 composite motifs in SOX2-dTAG mESCs. **D.)** Sampled single molecules underlying the averages shown as lineplots for each cluster. Each line represents 1,500 nucleotides extracted from an individual molecule, centered at the SOX2-OCT4 motif. **E.)** Heatmap representation of enrichment (red) or depletion (blue) of clusters comparing fibers at bound vs. unbound SOX2-OCT4 composite motifs. Data are shown for dTAG^V-^1-NEG- (designated as ‘NEG) and dTAG^V-^1-treated (time of treatment indicated) SOX2-dTAG mESC samples, stratified by biological replicate. Black dots mark Fisher’s exact tests where *q* > 0.1 (not significant). **F.)** Heatmap of Euclidian distances computed from enrichment data in (E); replicates cluster by drug treatment, suggesting that loss of SOX2 protein is associated with a reproducible change in accessibility states at SOX2-OCT4 motif loci. **G.)** Analysis of individual Leiden-defined clusters of nucleosome types at SOX2-OCT4 composite motifs. **Top:** Histograms of footprint sizes on a per-cluster basis. Middle: Mean accessibility traces centered on footprint midpoints on a per-cluster basis. **Bottom:** ‘Horizon plots’ for subfootprint sizes versus distance from footprint midpoints to assess subfootprint enrichment at particular locations on a per-cluster basis. z-scores are capped such that the maximum shown in the heatmap are the 95^th^ percentile values within ±70 bp of the respective footprint midpoints. **H.)** As in (G), but for SOX2-associated translational positions. Accordingly, mean accessibility and ‘horizon plots’ are instead plotted relative to motif positions. **I.)** Distribution of Spearman’s *r* values for pairwise comparisons of cluster abundances for SOX2-OCT4 motif-associated distortion patterns in large-scale E14 mESC dataset. A minimum of *n* = 500 footprints was required in order to exclude lowly-sequenced libraries from this correlation analysis. J.) As in (I), but for SOX2-associated translational positions. **K.)** Heatmap of log_2_-transformed odds ratios (ORs) for assessing enrichment / depletion of different nucleosome types at bound vs. randomly sampled SOX2-OCT4 composite motifs in dTAG^V-^1-NEG-treated SOX2-dTAG mESCs. ORs designated with a grey dot indicate those from Fisher’s exact test that are not statistically significant (*i.e.* Storey *q*-value > 0.1). Correlation is computed relative to E14 mESC data (data from **Figure 4C**). **L.)** Heatmap of effect sizes (Δ) for nucleosome types at bound SOX2-OCT4 motifs in dTAG^V-^1-NEG vs. dTAG^V-^1-treated (for 2 and 6 hr) SOX2-dTAG mESCs, stratified by biological replicate. Effect sizes designated with a grey dot indicate those from Fisher’s exact test that are not statistically significant (*i.e.* Storey *q*-value > 0.1). **M.)** As in **(K)**, but for SOX2-associated translational positions.

**Extended Data Figure 6:**
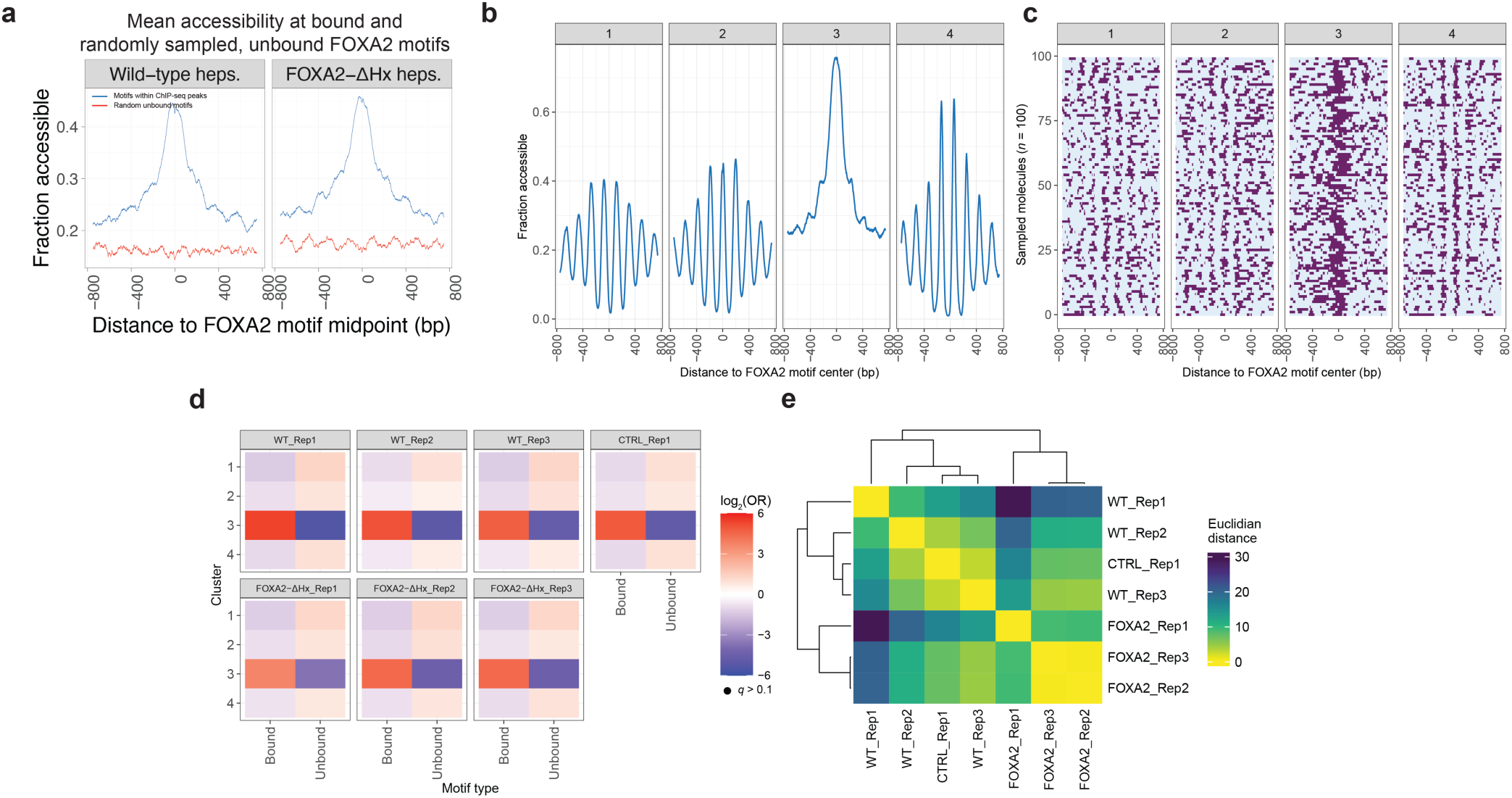
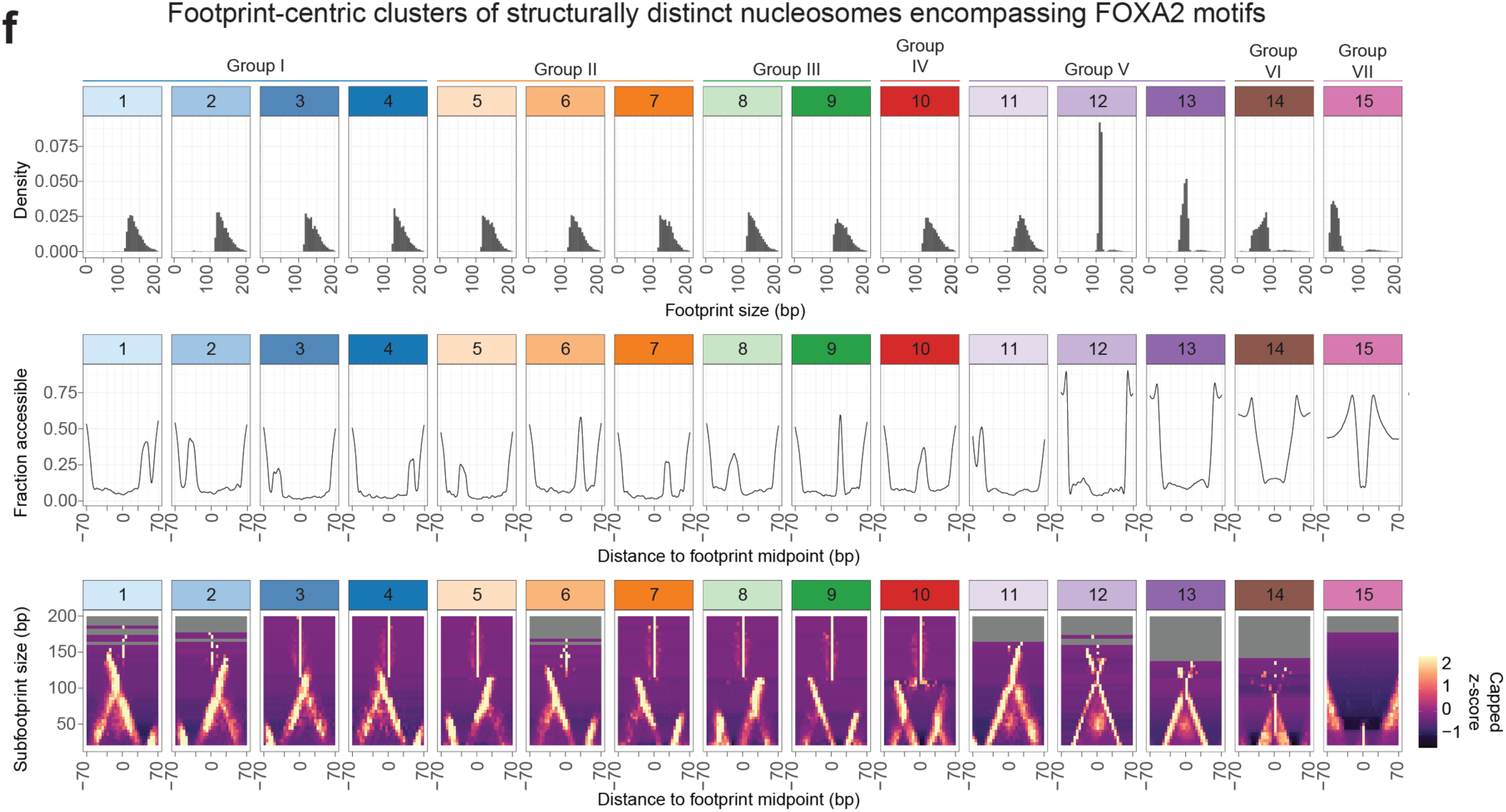
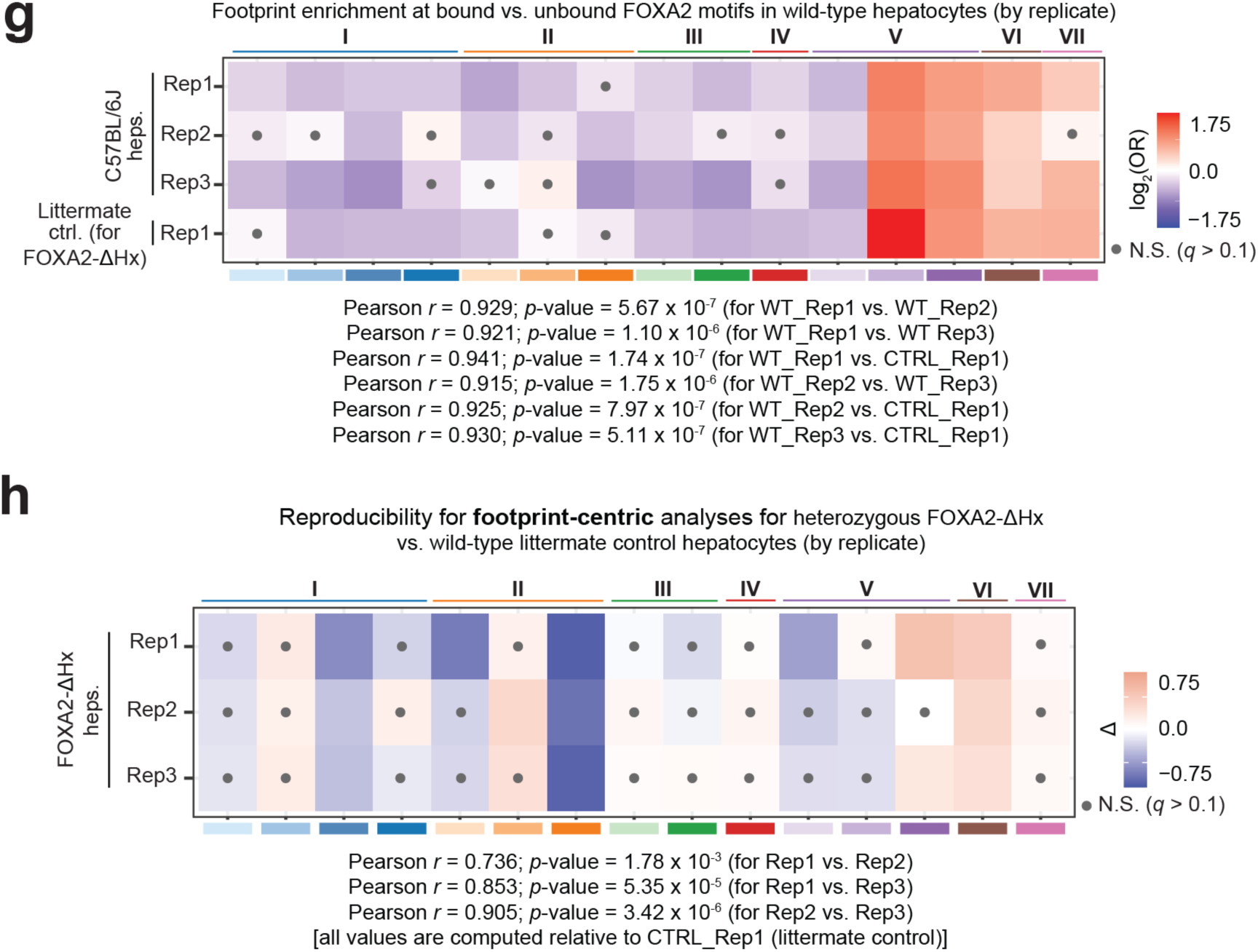
Validation and reproducibility of FOXA2-associated distortion patterns and translational positions, related to Figure 5. **A.)** Fraction accessible across ±750 bp window centered on bound (blue) and random (red) FOXA2 motifs. Comparison of data from wild-type and heterozygous FOXA2-ΔHx hepatocytes indicates that deletion of a FOXA2 histone-interacting domain is associated with minimal changes in mean accessibility at cognate FOXA2 binding sites. **B.)** Line plot representation of mean fraction accessible for Leiden-defined clusters at FOXA2 motifs in wild-type and heterozygous FOXA2-ΔHx hepatocytes. **C.)** Sampled single molecules underlying the averages shown as lineplots for each cluster. Each line represents 1,500 nucleotides extracted from an individual molecule, centered at the FOXA2 motif. **D.)** Heatmap representation of enrichment (red) or depletion (blue) of clusters comparing fibers at bound vs. unbound FOXA2 motifs. Data are shown for wild-type (samples designated as WT are from C57BL/6J mice; sample designated as CTRL serves as littermate control for FOXA2-ΔHx samples) and heterozygous FOXA2-ΔHx hepatocyte samples, stratified by biological replicate. Black dots mark Fisher’s exact tests where *q* > 0.1 (not significant). **E.)** Heatmap of Euclidian distances computed from enrichment data in (D); replicates cluster by genotype, suggesting that accessibility patterns are more similar within wild-type samples than to helical-domain-mutant samples, and vice versa. **F.)** Analysis of individual Leiden-defined clusters of nucleosome types at FOXA2 motifs. **Top:** Histograms of footprint sizes on a per-cluster basis. **Middle:** Mean accessibility traces centered on footprint midpoints on a per-cluster basis. Bottom: ‘Horizon plots’ for subfootprint sizes versus distance from footprint midpoints to assess subfootprint enrichment at particular locations on a per-cluster basis. z-scores are capped such that the maximum shown in the heatmap are the 95^th^ percentile values within ±70 bp of the respective footprint midpoints. **G.)** Heatmap of log_2_-transformed odds ratios (ORs) for assessing enrichment / depletion of different nucleosome types at bound vs. randomly sampled FOXA2 motifs in wild-type hepatocytes, stratified by biological replicate. ORs designated with a grey dot indicate those from Fisher’s exact test that are not statistically significant (*i.e.* Storey *q*-value > 0.1). **H.)** Heatmap of effect sizes (Δ) for nucleosome types at bound FOXA2 motifs in heterozygous FOXA2-ΔHx hepatocytes vs. littermate control, stratified by biological replicate. Effect sizes designated with a grey dot indicate those from Fisher’s exact test that are not statistically significant (*i.e.* Storey *q*-value > 0.1).

**Extended Data Figure 7:**
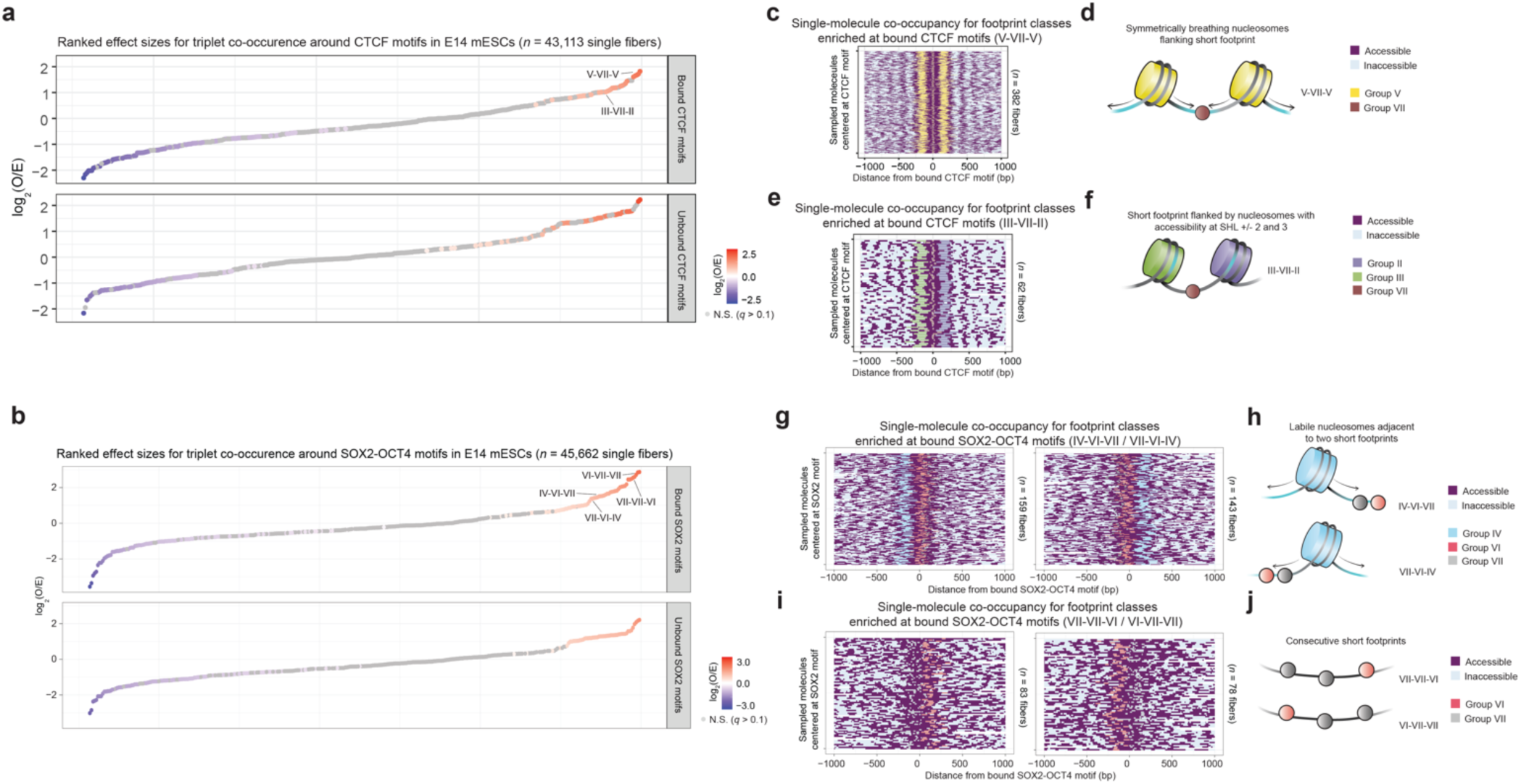
Co-occupancy analysis at CTCF and SOX2-OCT4 composite motifs, related to Figure 3-4. **A.)** Waterfall plots of ranked effect sizes for all possible ‘triplets’ at bound and random CTCF motifs. Grey dots used to indicate effect sizes for ‘triplets’ with *q* > 0.1. Red and blue dots are used to indicate ‘triplets’ that are enriched and depleted within a given locus, respectively. Specific ‘triplets’ enriched at bound, but not at unbound, CTCF motifs used for single-molecule visualization are indicated. **B.)** As in **(A)**, but for ‘triplets’ at bound and random SOX2-OCT4 composite motifs. **C.)** Single-molecule visualization of bound CTCF-associated chromatin fibers with V-VII-V ‘triplets’, wherein each row represents a unique sequenced fiber (total number of molecules indicated on right side of each plot). Accessibility patterns across a 2-kb window centered on domain midpoints of interest are also depicted (purple for EcoGII-accessible and teal for EcoGII-inaccessible DNA, respectively). Footprints from different groups are colored as described in associated legend; relative widths depict the number of nucleotides protected per footprint. **D.)** Schematic for ‘triplets’ with symmetrically breathing nucleosomes flanking short footprint (V-VII-V), enriched at bound CTCF motifs. **E-F.)** As in **(C-D)**, but for ‘triplets’ comprising short footprint flanked by nucleosomes with accessibility at SHL ±2 and ±3 (III-VI-II), enriched at bound CTCF motifs. **G-H.)** As in **(C-D)**, but for ‘triplets’ comprising labile nucleosomes adjacent to two shorter footprints (IV-VI-VII / VII-VI-IV), enriched at bound SOX2-OCT4 motifs. **I-J.)** As in **(C-D)**, but for ‘triplets’ comprising three consecutive short footprints (VII-VII-VI / VI-VII-VII), enriched at bound SOX2-OCT4 motifs.

## METHODS

### SAMOSA from E14 mESCs

We compiled a large compendium of SAMOSA data from wild-type E14 mESCs, comprising both newly generated datasets and published datasets from our laboratory, including: (1) our study on the function of ISWI-family remodeling complexes^29^; (2) samples prepared with our SMRT-Tag protocol, which utilizes a transposase-based strategy for PacBio library preparation from lower input amounts of footprinted DNA^27^; and (3) mESCs labeled with the halogenated thymidine analog, BrdU, for varying timepoints in our study of single-fiber accessibility patterns in newly-replicated chromatin^33^. Only samples with a mononucleosome-dinucleosome ratio (MDR; calculation detailed below in “Data analysis and visualization”) > 10 were included for analysis.

#### Cell culture

For newly generated libraries, E14 mESCs were grown on gelatin-coated (0.1% solution in 1x PBS) tissue culture plates. Cells were cultured in mESC media (DMEM with GlutaMAX (ThermoFisher 10566-016)), supplemented with: 15% FBS (ThermoFisher SH30071.03), 14.2 mM 2-Mercaptoethanol (Bio-Rad 1610710), 1x NEAA (ThermoFisher 11140-50), 1 mM sodium pyruvate (ThermoFisher 11360-070), and 1x LIF (purified by the laboratory of Barbara Panning (University of California, San Francisco)). Some samples were cultured with MEK (1 µM PD0325901 (Apex Bio / Fischer A3013-25)) and GSK3β (3 µM CHIR99021 (Apex Bio / Fischer A3011-100)) inhibitors (2i). Media was changed daily and cells were passaged with 1x TrypLE (ThermoFisher 12605010) when confluent. To harvest mESCs, cells were rinsed with 1x PBS (ThermoFisher 10010023), dissociated with 1x TrypLE, quenched with mESC media, and centrifuged at 500 g for 5 min. All mESC lines used in this study were regularly tested for mycoplasm contamination (Lonza LT07-318).

#### Methyltransferase footprinting and library preparation

Live cell counts were estimated (Countess 3) to ensure equal numbers of nuclei per unit methyltransferase were used in each footprinting reaction. Nuclei were isolated by incubating 1 x 10^6^ live mESCs in 1 mL ice-cold NE1 Buffer (20 mM HEPES, 10 mM KCl, 1 mM MgCl_2_, 0.1% Triton X-100, 20% glycerol, 1x Protease Inhibitor (Roche 4693132001)) on ice for 5 min. Nuclei were pelleted by centrifugation at 500 g for 5 min at 4°C. Nuclei were rinsed in 1 mL Buffer M (15 mM Tris-HCl pH 8.0, 15 mM NaCl, 60 mM KCl, 0.5 mM spermidine) and re-pelleted by centrifugation at 500 g for 5 min at 4°C. For footprinting, nuclei were resuspended in 200 µL Buffer M with 1 mM SAM (NEB B9003S) and 10 µL high-concentration EcoGII (custom order from NEB). Nuclei were incubated at 37°C for 30 min, with a spike-in of 1 µL 32 mM SAM stock halfway through the footprinting reaction. For light MNase digest to liberate chromatin fragments, footprinted nuclei were pelleted by centrifugation at 500 g for 5 min at 4°C, resuspended in 200 µL Buffer M with 1 mM CaCl_2_ and 0.02 U MNase (Sigma N3755), and incubated on ice for 30 min. To quench the MNase reaction, 2 mM EGTA was added.

To purify DNA, samples were treated with 10 µL RNaseA (ThermoFisher AM2270) for 10 min at 37°C, followed by the addition of 2 µL 10% SDS and 2 µL 20 mg/mL Proteinase K (ThermoFisher AM2548) for >2 hr (up to overnight) at 65°C. To extract DNA, an equal volume (∼250 µL) phenol-chloroform-isoamyl alcohol was added, samples were mixed by brief vortexing, and samples were centrifuged at max speed for 2 min. The aqueous phase was transferred to a fresh tube along with 0.1x volumes 3M NaOAc, 2x volumes ice-cold 100% ethanol, and 1 µL GlycoBlue co-precipitant (Thermo AM9516). Samples were mixed by inverting tubes and incubated overnight at -20°C, centrifuged at max speed for 30 min at 4°C, washed in 500 µL ice-cold 70% ethanol, air dried, and resuspended in 30 µL Buffer EB. DNA concentrations were measured by Qubit dsDNA High Sensitivity Quantification Kit (ThermoFisher Q32851). Up to 1 µg DNA was prepared into PacBio SMRT libraries with the SMRTbell Express Template Prep Kit 2.0 per manufacturer’s directions. Fragment size distributions were estimated by Femto Pulse (Agilent). Libraries were sequenced on PacBio Sequel II 8M SMRTcells.

### SAMOSA from SOX2-dTAG mESCs

Degron-tagged *Sox2^FKBP-F36V^* (hereafter referred to as SOX2-dTAG) mESCs were provided by the laboratory of Elzo de Wit (Netherlands Cancer Institute)^54^. SOX2-dTAG mESCs were cultured in the same mESC media described above (including 2i) and were plated at a density of 5 x 10^5^ per well in 6-well plates at 24 hr before harvesting. For degradation, SOX2-dTAG mESCs were treated with 0.5 µM dTAG^V^-1 (Tocris 6914) for varying timepoints (30 min, 2 hr, or 6 hr). As control, SOX2-dTAG mESC samples were treated with 0.5 µM dTAG^V^-1-NEG (Tocris 6915) for 6 hr before harvesting. All experiments were performed in biological duplicate. Footprinting and downstream steps were performed as detailed in the section above, except in lieu of the MNase digest step, we sheared purified genomic DNA with Covaris g-TUBEs (Covaris 520079; 4,600 g for 6 passes).

#### Western blot for SOX2 protein in nuclear extracts

To validate levels of SOX2 knockdown, mESC nuclei were isolated in NE1 Buffer, as done for matched methyltransferase-footprinted samples. Pelleted nuclei were resuspended in 100 µL ice-cold RIPA buffer (150 mM NaCl, 1% NP-40, 0.5% sodium deoxycholate, 0.1% SDS, 50 mM Tris pH 7.4, 1x Protease Inhibitor (Roche 4693132001)). Samples were incubated for 30 min on ice and sonicated briefly to solubilize chromatin. Protein extracts were clarified by centrifugation at 12,000 g for 20 min at 4°C. Soluble material was collected, protein concentrations were measured by BCA (ThermoFisher 23227), and samples were diluted in NuPAGE LDS Sample Buffer (ThermoFisher NP0007) with 5% BME. Samples were reduced by boiling for 10 min at 95°C, resolved on 4-12% Bis-Tris gels (ThermoFisher NP0322), and transferred to PVDF membranes. Membranes were incubated overnight with the following primary antibodies: anti-SOX2 (CST 23064S; 1:1,000 dilution) and anti-Vinculin (Sigma V9264; 1:1,000 dilution; used as loading control). After washing, membranes were incubated with secondary antibodies conjugated to IRdye 700 or 800 (1:10,000 dilution) and imaged with the LiCor Odyssey.

### SAMOSA from CTCF-AID mESCs

CTCF-AID mESCs (EN52.9.1) were generated by the laboratory of Elphège Nora (University of California, San Francisco)^47^. Raw data from CTCF-AID samples used in this study were previously published by our group^33^, and samples were required to have a mononucleosome-dinucleosome ratio (MDR; calculation detailed below) > 10 to be included in analyses. This criteria resulted in: (1) *n* = 10 biological replicates of auxin-treated (for 6 hr) CTCF-AID mESCs; and (2) *n* = 6 biological replicates of DMSO-treated CTCF-AID mESCs (as controls).

### SAMOSA from primary mouse hepatocytes

#### Mouse husbandry and genotyping

All animal experiments were performed under the supervision and approval of the Institutional Animal Care and Uses Committee (IACUC) and the University of California, San Francisco (Protocol # AN179718-03F). 6-week old, C57BL/6J mice (Jackson Laboratory #000664) were used as wild-type hepatocyte samples in our dataset (*n* = 3 biological replicates). *Foxa2*^ΔHx-*TagRFP*^-expressing mice (hereafter referred to as FOXA2-ΔHx) were provided by the laboratory of Kenneth Zaret (University of Pennsylvania)^58^. Samples in this study included heterozygous FOXA2-ΔHx mice (*n* = 3; 1 male and 2 females) and a littermate wild-type control male mouse.

Genotyping for the presence (∼230 bp) or absence (∼200 bp) of the associated Tag-RFP fluorophore was performed with the following primer pair:

lox2272_F: AGTGTTGTCTTCTGCCTTTGAG lox2272_R: GCTTACCTTAGTCTCGGTCTTGG

Genotyping for the presence (∼300 bp) or absence (∼270 bp) of the helical domain in FOXA2 was performed with the following primer pair:

TagRFP_F: GCTCTTCGCCCTTAGACACC TagRFP_R: ATCAGCCCCACAAAATGGAC

#### Isolation of primary mouse hepatocytes

Hepatocyte isolation was performed by the Liver Cell Isolation, Analysis, and Immunology Core of the UCSF Liver Center. After flushing the liver free of blood, the organ was perfused for 3 min at 4 mL/min with Liver Perfusion Medium (ThermoFisher 17701038), followed by an additional 7 min at 4 mL/min with Liver Digest Medium (ThermoFisher 17703034). The liver was then removed, diced with scissors, and suspended in DME-H21 with insulin-transferrin-selenium (ThermoFisher 41400045), penicillin-streptomycin, and 5% FBS. The crude suspension was strained through a 100-µm filter and pelleted twice at 60 g with interval washing. Hepatocytes were purified by suspension of the pellet and centrifugation through 45% Percoll (Cytiva 17089102) at 130 g for 15 min. Purified hepatocytes, which pelleted through the Percoll, were suspended in DME-H21 as above. Final hepatocyte suspensions had a viability > 90%.

#### Methyltransferase footprinting for hepatocyte samples

Footprinting reaction for samples denoted HepWT_Rep1 and HepWT_Rep2 was performed in 1 x 10^6^ purified hepatocyte nuclei per library. Footprinting and downstream steps were performed as described above, except primary hepatocytes were initially dounced in NE1 Buffer (10x with loose pestle, 10x with tight pestle) to facilitate nuclear isolation.

The footprinting reaction for all remaining hepatocyte samples was performed with 1 x 10^6^ digitonin-permeabilized hepatocytes. Footprinting and downstream steps were performed as described above, with two notable exceptions. First, in lieu of isolating nuclei with NE1 Buffer, we directly permeabilized cells by adding 0.05% digitonin (ThermoFisher BN2006) to Buffer M when performing the EcoGII footprinting reaction. Second, instead of treating nuclei with MNase to liberate chromatin fragments, we sonicated purified genomic DNA with the Megaruptor 3 (concentration = 10 ng/µL, volume = 100 µL, speed = 031) to achieve more consistently-sized and longer molecules (>10 kb).

### Preprocessing PacBio data for single-molecule accessibility

To preprocess PacBio data, we utilized software from Pacific Biosciences and a custom script (*hmm_output_t_values.py*) to successively run our NN-HMM^29^ across a range of user-specified transition probabilities. These resulted in the following output per sample: (1) an alignment of CCS reads to the mouse reference genome (mm10); (2) an accessibility prediction per CCS molecule (Viterbi path of HMM component of model); and (3) an atlas of footprint locations (*i.e.* start / end positions) and sizes on a per-molecule basis. Files for (2) and (3) were generated for *t* = 1, 31, 51, 71, and 101 / 1,000 for each sequencing library.

### Data analysis and visualization

#### Quantifying extent of methyltransferase footprinting per sequencing library

We utilized the ratio of mononucleosome-sized to dinucleosome-sized footprints (MDR) as a proxy for measuring the extent of EcoGII footprinting on a per-sample basis. Poorly methylated samples exhibit a higher fraction of footprints longer than the expected length of DNA protected by a mononucleosome (*i.e.* lower MDR values), which can be ascribed to the failure of EcoGII to efficiently methylate in linker DNA. We employed a custom script (*compute_mdr_per_sample.py*), which uses the *find_peaks* function (from *scipy*) and computes the ratio of maximum peak heights in footprint length histograms for mononucleosome- and dinucleosome-sized footprints per sequencing library.

#### Visualizing single-molecule data across different t-values at single genomic loci

To visualize SAMOSA data at loci of interest, data were processed with a custom script (*generate_bam_for_igv.py*) to encode single-molecule accessibility patterns (for any given *t*-value) as an additional flag in aligned *.bam* files. These *.bam* files were concatenated across E14 mESC samples with MDR > 10 and visualized at the *Sox2* locus (chr3: 34,756,929 – 34,759,161; including the *Sox2* promoter, gene body, and downstream SCR) using a custom version of IGV (https://github.com/RamaniLab/SMRT-Tag/tree/main/igv-vis).

Bulk ATAC-seq data from mESCs were obtained as processed *.bw* files (GSE98390) and visualized at identical coordinates on the UCSC Genome Browser.

#### Identifying fibers that overlap annotated epigenomic domains in mESCs

A custom script from our group (https://github.com/RamaniLab/SAMOSA-ChAAT/blob/main/scripts/zmw_selector.py) was adapted to identify sequencing reads where a portion of the read falls within ±1 kb of midpoints of histone post-translational modification-defined epigenomic domains in mESCs (*i.e.* H3K4me1, H3K4me3, H3K9me3, H3K27me3, and H3K36me3 ChIP-seq peaks; derived from ENCODE data).

#### Identifying fibers that overlap repeat elements in mouse genome

A custom script from our group (https://github.com/RamaniLab/SAMOSA-ChAAT/blob/main/scripts/blast_bam.sh) was used to identify sequencing reads with one or more matches to repeat elements of interest. In brief, we ran BLAST on CCS reads from a database consisting of mouse major satellite sequence (M32564.1), minor satellite sequence (X14462.1), and telomeric DNA sequence (pentameric repeats of TTAGGG). For fibers with multiple repeat matches, only the repeat sequence with the lowest E-value was considered for downstream analyses.

#### Identifying fibers that overlap bound and unbound transcription factor motif matches

To define instances of TF-bound motifs, we referenced the following datasets: ChIP-nexus data for SOX2 in mESCs (GSE137193)^53^, ChIP-seq data for CTCF in mESCs (ENCFF508CKL), and ChIP-seq data for FOXA2 in mouse hepatocytes (GSE157452)^59^. For SOX2 motifs, we retained only the subset of peaks that contained a SOX2-OCT4 composite motif match as defined by this dataset^68^. For CTCF data, we utilized a curated set of ChIP-seq backed motifs initially processed in this study^45^. For FOXA2 data, we retained only the subset of peaks that contained a PWM match (MA0047.2) for the FOXA2 motif^69^. For control sites, we randomly sampled an equal number of motif matches per factor of interest from a previously published list of motifs across the mouse genome^68^, requiring that these motifs fall at least 1 kb away from the nearest ChIP-annotated peak. As above with epigenomic doamins, we utilized a custom script to identify sequencing reads where a portion of the read falls within ±1 kb of motif midpoints.

#### Clustering single-molecule accessibility patterns with respect to footprint midpoints

To examine patterns of nucleosomal distortion, we clustered accessibility patterns across a range of *t*-values with respect to footprint midpoints. Footprints <200 nt in length (*i.e.* intended to capture both nucleosomal and subnucleosomal species) within ±500 bp of midpoints of epigenomic domains, repeat sequences, or TF-binding motifs were selected. On a per-footprint basis, accessibility data (from *t* = 31, 51, and 71 / 1,000) for a 140-bp window centered on each footprint midpoint (*i.e.* 420 bp in total per footprint) was used as input for Leiden clustering (*res* = 0.5, 0.6, 0,6, and 0.6 for domains / repeats, SOX2, CTCF, and FOXA2 motifs, respectively). Clusters were filtered such that the least abundant clusters that collectively summed up to <10% of each dataset were removed. It should be noted that: (1) accessibility data was oriented accounting for sequence feature strand before clustering to preserve any potential directional effects associated with non-palindromic TF-binding motifs; (2) this approach was intended to enable clustering on accessibility information across multiple *t*-values per footprint; and (3) this window size was selected to prioritize signal from SHL -7 to SHL +7 for nucleosomal footprints. Representative code for performing this analysis is provided as a custom script (*process_footprints_nucleosomal_distortion.py*).

#### Clustering single-molecule accessibility patterns with respect to sequence features of interest

To define translational settings, we clustered accessibility patterns across a range of *t*-values with respect to midpoints of a given sequence feature of interest. Footprints <200 nt in length wherein the absolute distance between the footprint midpoint and sequence feature was less than half the total length of the footprint in question (*e.g.* within ±100 bp for a 200 nt footprint) were selected. On a per-footprint basis, accessibility data (from *t* = 31, 51, and 71 / 1,000) for a 140-bp window centered on each sequence feature (*i.e.* 420 bp in total per footprint) was used as input for Leiden clustering (*res* = 0.3, 0.8, 0.8, and 0.8 for repeats, SOX2, CTCF, and FOXA2 motifs, respectively). Clusters were filtered such that the least abundant clusters that collectively summed up to <10% of each dataset were removed. Representative code for performing this analysis is provided as a custom script (*process_footprints_translational_positions.py*).

#### Plotting footprint size distributions, mean accessibility patterns, and ‘horizon plots’ per cluster

To assign putative biological function to clusters of nucleosomal distortion patterns or translational positions, we visualized the following features. First, we plotted the distribution of footprint sizes on a per-cluster basis for data from the lowest *t*-value examined (*t* = 1 / 1,000). Second, we plotted mean accessibility as a function of distance to respective footprint / sequence feature midpoints from the highest *t*-value used for Leiden clustering (*t* = 71 / 1,000). Third, we generated ‘horizon plots’ (analogous to commonly used ‘V plots’ in MNase-seq data) by computing z-scores for enrichment of footprints of particular size (20 – 200 nt) at specific positions (±70 bp from footprint / sequence feature midpoints) for each individual cluster. z-scores are summed per size-position combination across the range of *t*-values previously used for clustering (*t* = 31, 51, and 71 / 1,000). Color scale limits for the resultant heatmap are capped by the 0th (lower) and 95th (upper) percentile z-scores for positions within ±70 bp of the footprint / sequence feature midpoint. Clusters of entirely unmethylated footprints were manually filtered out before downstream quantification. Representative code for performing this analysis is provided as a custom script (*plot_footlen_meanacc_horizondata.py*).

#### Identifying groups of clusters with shared nucleosomal distortion patterns / translational positions

To group Leiden assignments for nucleosomal distortion patterns or translational positions in an unbiased manner, we performed hierarchical clustering of ‘horizon plot’ data as follows. z-scores per cluster were filtered such that only those corresponding to footprints <100 nt and <50 nt in length were retained for defining nucleosomal distortion and translational position groups, respectively. This choice was intended to prioritize clustering of ‘horizon plot’ data specifically within subfootprint size ranges that capture subnucleosomal organization. Data were manually inspected and z-scores flipped with respect to footprint / sequence feature midpoints to account for the nucleosomal dyad as an axis of symmetry. The distance matrix was computed with the *dist* function (in R), hierarchical clustering was performed with *hclust* (from *factoextra* package; *method* = ward.D2), and output was visualized as a dendrogram (with *fviz_dend*).

#### UMAP visualization of footprint types within epigenomic domains and at repeat sequences

From accessibility data for footprints within histone modification-defined domains and at mouse repeat elements, we employed Scanpy (v1.9.3) for PCA-based dimensionality reduction, construction of a *k*-nearest neighbors graph (*metric* = correlation, *n_neighbors* = 15), and UMAP visualization (*min_dist* = 0.5). Clusters were colored according to Leiden assignments and proportions of each Leiden-defined cluster were represented as a pie chart surrounding the UMAP visualization.

#### Structural visualization of nucleosomal accessibility patterns

For structural visualization of mean accessibility patterns from defined clusters, we utilized a custom script (*accessibility_to_pdb_structure.py*) to convert our mean single-molecule accessibility data per cluster of interest from the highest *t*-value used for Leiden clustering (*t* = 71 / 1,000) into a format compatible with visualization via ChimeraX (v1.7.1; *attribute* = percentAcc, *match mode* = 1-to-1, and *recipient* = residues)^70^. Accessibility values were overlaid on the following PDB structure: 7KBD^71^.

*Computing enrichment of nucleosomal distortion patterns / translational positions across distinct genomic loci* To compute odds-ratios for Leiden-defined clusters across epigenomic domains and repeat elements, or for bound vs. randomly-sampled TF motifs, Fisher’s exact tests were carried out with *scipy* (in Python). All computed *p*-values were corrected with a Storey *q*-value correction, using the *qvalue* package (in R).

#### Computing effect sizes associated with TF knockdown or across genotypes

To compute effect sizes upon degron-mediated TF knockdown or across mouse genotypes, Fisher’s exact tests were carried out for footprints specifically at TF-bound motifs (*e.g.* footprints from dTAG-^V^1 vs. DMSO-treated SOX2-dTAG mESCs) with *scipy* (in Python). All computed *p*-values were corrected with a Storey *q*-value correction, using the *qvalue* package (in R).

#### Reproducibility of nucleosomal distortion patterns or translational positions across replicates

To assess reproducibility in our large-scale E14 mESC dataset, we asked if the rank-order of clusters of nucleosomal distortion patterns or translational positions is preserved across sequencing libraries. To do this, the relative proportion of each Leiden-defined cluster at domains / repeats or at TF-bound motifs was computed on a per-sample basis and analyzed using Spearman’s correlation (with *spearmanr* from *scipy*). Footprints from all sequencing libraries with MDR > 10 were included in Leiden clustering of accessibility patterns; this choice was intended to include as many footprints as possible to increase statistical power. However, for these correlation analyses, we imposed a minimum sequencing depth for a biological replicate to be included, with the goal of assessing reproducibility in sufficiently-sequenced samples (minimum number of footprints indicated on each respective plot).

To evaluate reproducibility in SOX2-dTAG mESCs, CTCF-AID mESCs, and FOXA2-ΔHx hepatocytes, we calculated the Pearson’s correlation (with *pearsonr* from *scipy*) for effect sizes computed on a per-sample basis. This choice was intended to capture potential quantitative changes in cluster proportions between degron conditions / genotypes across replicates.

#### Identifying single-molecule co-occupancy patterns for different footprint groups

Individual fibers with at least three Leiden-classified footprints within ±500 bp of repeat elements / TF motifs of interest were selected. Fibers with footprints >200 nt and/or with footprints in the least abundant 10% of clusters were excluded from further analysis. Consecutive ‘triplet’ footprints were tabulated by computing midpoints and lengths for: (1) the central-most footprint nearest the repeat element / TF-binding motif; (2) the most proximal footprint upstream of the central-most footprint; and (3) the most proximal footprint downstream of the central-most footprint. Per-molecule accessibility data was computed for a 2-kb window centered on the repeat element / TF-binding motif of interest for visualization alongside footprint positions, accounting for sequence feature strand appropriately. Representative code for performing this analysis is provided (*process_triplet_footprints.py*).

#### Computing enrichment of consecutive ‘triplets’ on single chromatin fibers

Individual clusters of nucleosomal distortion patterns were combined based on groups defined by shared subfootprint organization from hierarchical clustering, as described above. Expected frequencies of footprint ‘triplets’ were computed as the product of the independent probabilities of observing each group within a given dataset. These values were used to calculate log_2_(O/E) ratios for each ‘triplet’ across epigenomic domains, repeat elements, or TF-binding motifs. Enrichment was assessed using Fisher’s exact test (*fisher.test* function in R), with FDR correction applied for multiple comparisons. The results were visualized as a ranked-order scatterplot, reflecting the effect size for each ‘triplet’.

#### Visualizing mean accessibility patterns at bound vs. random transcription factor motifs

Previously-published scripts from our group were used to compute and cluster mean accessibility observed within ±750 bp of TF-binding motifs across different conditions (*i.e.* before / after TF knockdown; across genotypes)^33^. Euclidian distances for resultant odds-ratios for bound vs. unbound motifs per sample are plotted to visualize their similarity across conditions and replicates.

## DATA AND CODE AVAILABILITY

Raw and processed data will be made available at GEO accession GSEXXXXXX. Custom scripts used for analyses are available at https://github.com/RamaniLab/IDLI.

## ACKNOWLEDGEMENTS

We thank members of the Ramani Lab, Siva Kasinathan (Stanford), Hiten Madhani (UCSF), Geeta Narlikar (UCSF), and Srinivas Ramachandran (University of Colorado) for helpful discussions and comments on the manuscript. We acknowledge Kenneth Zaret (University of Pennsylvania) for sharing Foxa2-ΔHx mice, Jackson Goudreau and Michael Mui for assistance with mouse husbandry, Tami Tolpa for generating schematics used in our figures, and support from core facilities within the UCSF Liver Center (P30-DK026743). MGY was supported by NIH T32-DK060414 and by the Gladstone-CIRM Scholars Program. This work was partially supported by NIH grants U01-DK127421 and DP2-HG012442 to VR. VR acknowledges generous support from the Searle Scholars Award and the W.M. Keck Foundation.

